# Hidden Markov Models based search in combination with structural bioinformatics pipeline leads to the identification of DAF-12 distant orthologous in *Meloidogyne incognita*

**DOI:** 10.1101/2023.07.19.549723

**Authors:** Claudio David Schuster, Vanessa Judith Santillán Zabala, Rafael Betanzos San Juan, Ezequiel Sosa, Cristian Rodriguez, Maria Florencia Kronberg, Eliana Rosa Munarriz, Gerardo Burton, Olga Alejandra Castro, Carlos Pablo Modenutti

**Author notes:** These authors contributed equally to this work (Claudio David Schuster, Vanessa Judith Santillán Zabala, Rafael Betanzos San Juan).

## Abstract

Root-knot nematode (RKN) *Meloidogyne* spp. is one of the most damaging parasites due to its wide range of hosts. Here, we report a *C. elegans* receptor DAF-12 ortholog gene in *Meloidogyne incognita* (DAF-12_Minc_), a promising molecular target to modify the RKN life cycle. Using a combination of Hidden Markov Models (HMM) based sequence search and phylogenetic analysis we identified three DAF-12_Minc_ genes. Although the global sequence identity between previously reported DAF-12 genes and DAF-12_Minc_ was acceptable, the correlation between binding site residues was low in the multiple sequence alignment (MSA). Since those residues are critical for DAF-12 interaction with its ligand, the dafachronic acids (DAs), and thus its biological role, we investigated whether even if the sequence conservation is low, the active site structure was conserved and thus able to bind DAs. For this purpose, we built accurate homology models of DAF-12_Minc_ and used them to identify and characterize the ligand binding site (LBS) and its molecular interactions with DAs-like compounds. Finally, we cloned, expressed, and evaluated the biological role of DAF-12_Minc_ *in vitro* and *in vivo* using a DAF-12 antagonist. These *in vivo* results suggest that our strategy was effective to find orthologous genes among species even when sequence similarity is low.

**Author summary:** Root-knot nematodes are parasitic to plants and responsible for causing a significant loss of millions of dollars every year in crops worldwide, which makes it necessary to develop effective strategies to combat them. One popular approach is to identify genes that can serve as molecular targets. Typically, such molecular targets are discovered through basic research on model organisms. However, since they can be quite different from the target organism, conventional tools may not always be efficient in extrapolating results.

Dafachronic acids (DAs) are a crucial class of steroid hormones that regulate the development and physiology of nematodes. They are synthesized from cholesterol and are controlled by a nuclear hormone receptor known as DAF-12. This receptor acts as a master regulator of gene expression, playing a vital role in nematode biology, including development, reproduction, metabolism, stress response, and longevity. Therefore, DAF-12 is a promising molecular target for controlling parasitic nematodes.

Although DAF-12 was initially discovered in the model organism *Caenorhabditis elegans* and subsequently found in some parasitic nematodes, previous attempts to identify molecular targets in the *Meloidogyne* genus failed to detect DAF-12 orthologs. To address this gap, we employed a combination of sequence and structure analysis to identify potential candidates for DAF-12, a known and validated molecular target, which had not yet been found in *Meloidogyne incognita*. Our bioinformatics predictions were experimentally validated, which may serve as a starting point for future campaigns aimed at developing parasite control strategies based on this relevant molecular target.

## Introduction

Damages on crops produced by nematode infections are estimated to be valued at USD 157 billion annually [1]. In particular, the root-knot nematode (RKN) *Meloidogyne* spp. is one of the most damaging parasites due to its wide range of hosts [2]. The RKN second-stage juveniles (J2) penetrate behind the root tip and migrate intracellularly to reach the vascular cylinder where it induces the differentiation of 5 to 7 cells to establish its feeding site, which are known as giant cells. In most plant species, these giant cells are surrounded by hypertrophied cortical cells forming root-knot galls. These multinucleated giant cells provide nutrients to carry on the nematode life cycle. Soon after initiation of a feeding site, the J2 becomes sedentary and undergoes three molts to become an adult female that lays eggs into a gelatinous matrix in the root. After egg eclosion, J2 larvae locate root tips of suitable plant species and initiate a new cycle of infection [3][4][5]. An interesting aspect of J2 larvae is that it is a stage of vulnerability for the parasite, which makes it an attractive target to develop a controlling strategy [6].

Previous studies have proposed a wide variety of genes as potential molecular targets to fight nematode parasites, such as DAF-16 and SKN-1 [7]. Another potential interesting target is DAF-12, a nuclear receptor present in *Caenorhabditis elegans* that binds to a type of ligands known as dafachronic acids (DAs). DAs are cholesterol metabolites with an acidic carboxylic group in the side chain introduced by the cytochrome P450 oxidase DAF-9. In environmentally unfavorable conditions, DA synthesis is repressed and *C. elegans* arrests its development at an alternative stage called dauer larva. Upon returning to favorable conditions, DA synthesis is re-initiated, and the dauer larva exits the diapause and resumes its reproductive life cycle [8]. All of this indicates that DAF-12 plays a major role in regulating the *C. elegans* life cycle. Interestingly, several characteristics such as a sealed mouth, filariform shape, and thickened body wall cuticle, make the J2 *Meloidogyne* spp. larvae resemble the dauer larvae of the free-living nematode *C. elegans* [9].

A number of *C. elegans* DAF orthologous genes, including the DAF-12 nuclear receptor, have been identified in genomes of different parasitic nematodes species such as *Pristionchus pacificus, Bursaphelenchus xylophilus, Panagrellus redivivus*, *Ancylostoma celanicum, and Strongiloides stercoralis* [9]. The last two have crystallographic structures deposited in the Protein Data Bank [10] while *P. pacificus, B. xylophilus and P. redivivus* have not been structurally characterized. Based on these data, it is possible to hypothesize that the endocrine network mediated by the DAs that control the *C. elegans* life cycle may be also functional in *M. incognita*. Given the central role of DAF-12 in *C. elegans* life cycle, its ortholog gene in *M. incognita* could be a promising molecular target, but previous studies failed to find DAF-12 in *M. incognita* [9].

In this study we proposed a new strategy to find DAF-12 ortholog genes in *M. incognita*. This strategy takes into consideration not only gene and protein sequence similarity, but also protein domain structure using Hidden Markov Models (HMMs). Furthermore, we perform structural analyses to assess that the candidate proteins are able to bind the correspondent DA ligand. Using this combined strategy, we were able to successfully identify DAF-12 in *M. incognita*.

## Results

### Identification of orthologs of DAF-12 and DAF-9

In order to find possible orthologs of DAF-12 and DAF-9 in *M. incognita*, we performed a HMM based search over the *M. incognita* proteome [11] and identified 52 DAF-12-like genes (S6 File). Pairwise identity matrices (Fig 1) reveal that nucleotide sequences align overall better than the corresponding proteins.

**Fig 1.**
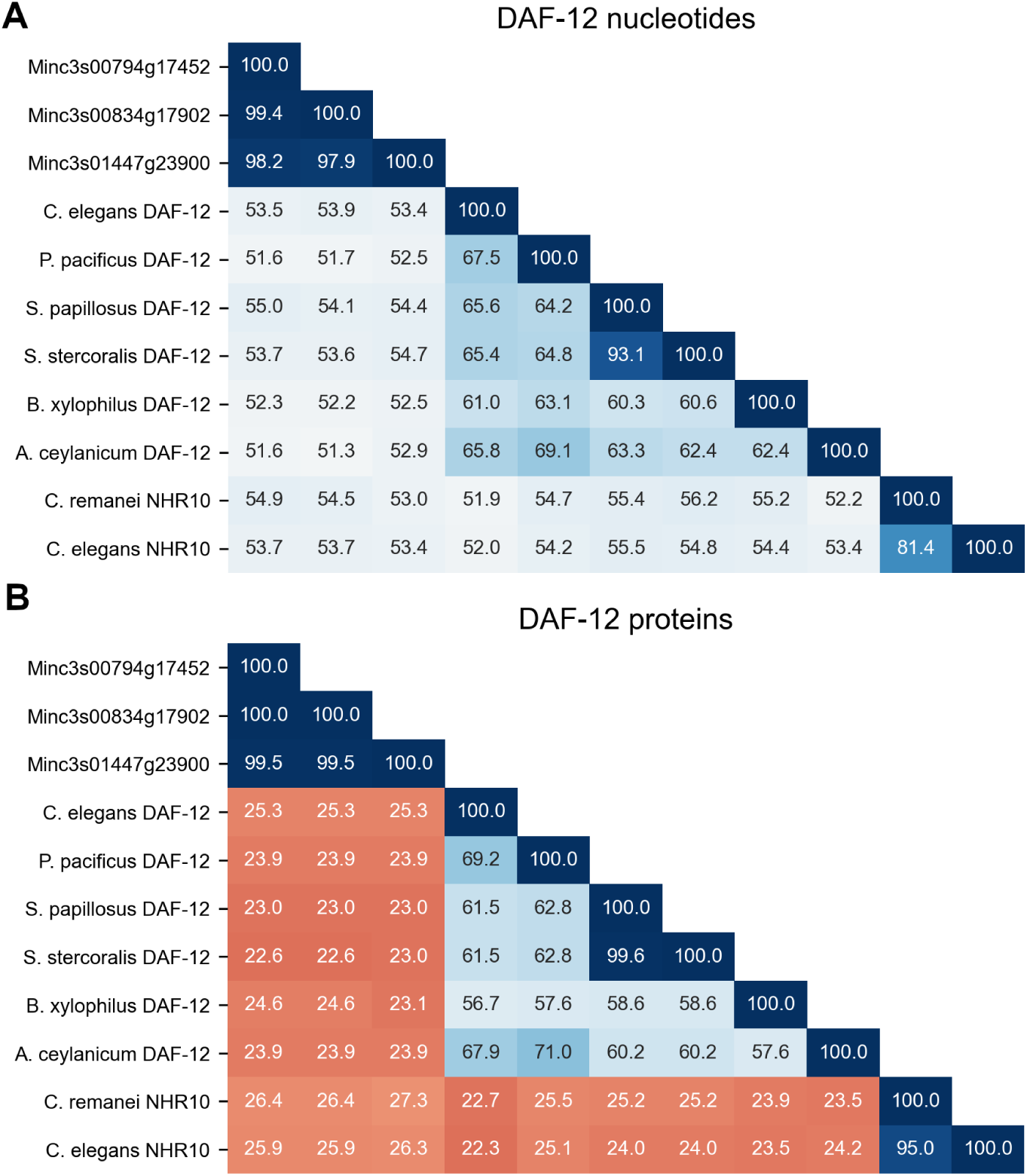
Pairwise identity matrices between *M. incognita* DAF-12 gene candidate sequences, reference sequences and outgroup sequences used in phylogenetic analysis. (A) Identity matrix between DAF-12 nucleotide sequences. (B) Identity matrix between DAF-12 protein sequences. Numbers indicate the percentage of pairwise identity between sequences. Color scale goes from lower sequence identity (red) to higher sequence identity (blue).

Phylogenetic analysis of DAF-12-like (Fig 2) genes and proteins showed potential orthologs in *M. incognita*, given that the same DAF-12-like genes grouped with the respective references in nucleotide and protein phylogenetic trees. Three *M. incognita* sequences (S10 Fig) were detected as strong candidates that display high identity percentages between themselves (Fig 1). Interestingly, the proposed orthologs still display quite low nucleotide and protein sequence identity when compared to those of other nematodes.

**Fig 2.**
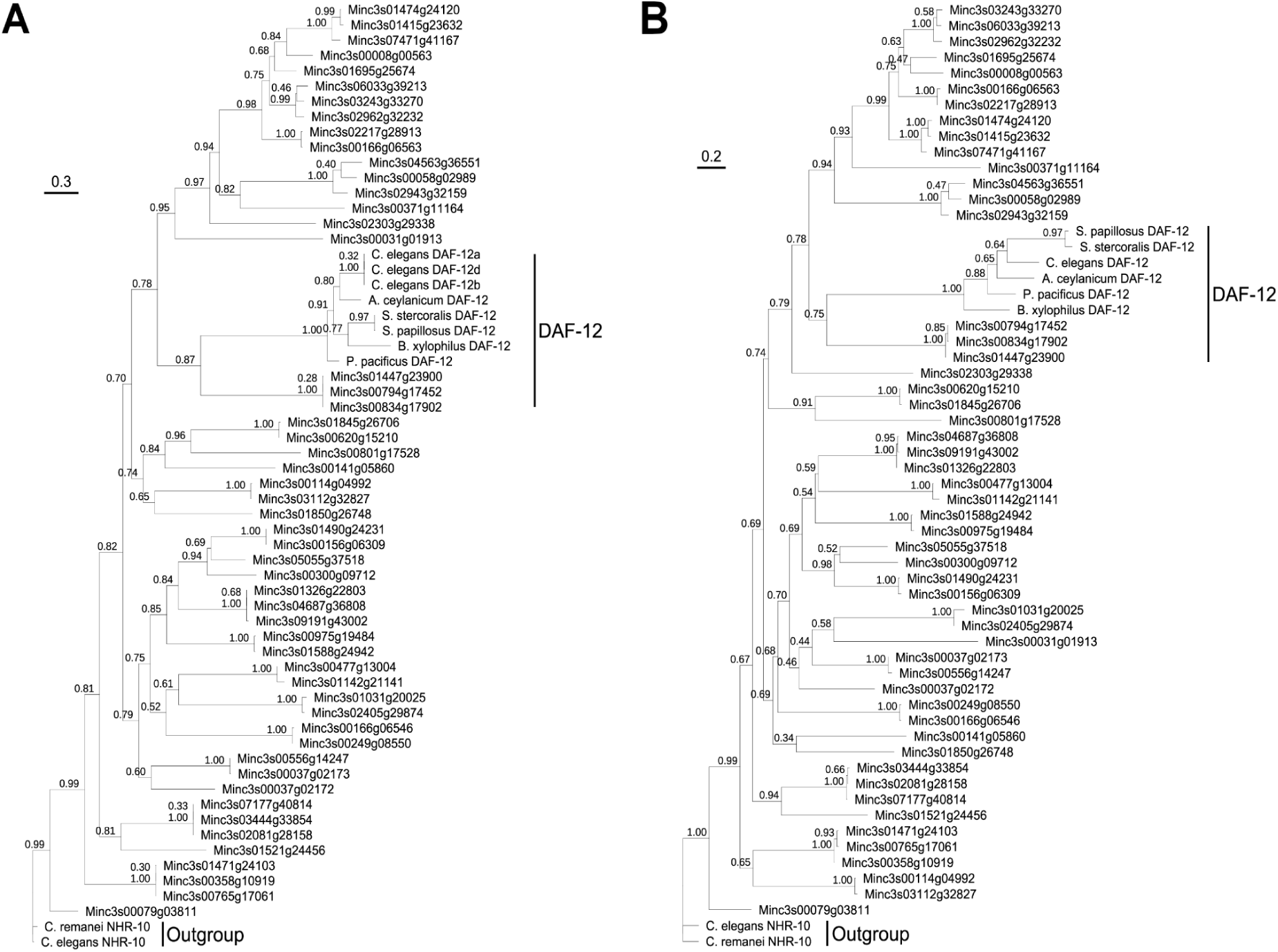
Phylogenetic trees inferred from *M. incognita* DAF-12-like nucleotide sequences (A) and protein sequences (B) obtained from *M. incognita* proteome data mining and DAF-12 reference sequences. Branch length is proportional to genetic change, tree scale shows genetic change per length unit, numbers on the branches indicate bootstrap values, clade label “DAF-12” comprises DAF-12 reference sequences and the 3 final candidates to DAF-12_Minc_.

From a general perspective, these results strongly suggest 3 potential DAF-12 orthologs in *M. incognita*. This narrowing down to three candidates is consistent with a previous genomic work on *M. incognita* that suggested it might be a triploid organism [11].

The same analysis was done to search for DAF-9, which performs the last step in DAs synthesis, and similar results were obtained (S7 File, S1, S2, S7 and S8 Figs).

### DAF-12 sequence analysis

Once the final three DAF-12 candidates in *M. incognita* were identified, the correspondent mRNA sequences were aligned to the respective genomic sequences of the most recent version of *M. incognita* genome as well as the genomic sequences of an older version of the genome (S9 Fig) [1]. The protein sequences of the three candidates were also aligned to hypothetical DAF-12 proteins from the predicted proteome of the old version of *M. incognita* genome, which were previously identified with Blast [12]. By comparing the sequences in both alignments three interesting observations emerge: (1) the candidate Minc3s00794g17452 differs significantly from the other two at the first 26 amino acids because the genomic sequence is shorter than the other two at the 5’ end (thus setting its starting codon in an intronic segment that is not translated in the other two candidates); (2) one of the predicted proteins (Minc10028) from the old genome version probably has been wrongly predicted, since it displays a segment unique to it that derives from translating an intronic fragment. In fact, an analysis performed with software Augustus [13] on the whole gene sequences of DAF-12 in the old genome version failed to correctly predict the proteins; (3) the other DAF-12 protein prediction from the old genome version (Minc18013) lacks 7 residues.

Based on these MSAs we decided to perform structural, *in vitro* and *in vivo* analyses on the sequence Minc3s00834g17902.

### Structural modeling of DAF-12 candidates

Although phylogenetic analysis suggested that the most likely DAF-12 orthologs were Minc3s00794g17452, Minc3s00834g17902 and Minc3s01447g23900, we observed low correlation in the (MSA) between the position of amino acids of the binding site in *M. incognita* and the previously reported sequences (S10 Fig). Considering that a single genetic mutation can trigger a cascade of compensatory mutations to preserve protein conformational and functional stability, we explored whether the ability to bind DA ligand was present in the structure, despite lack of specific active residue conservation.

To this end, we developed a structural bioinformatics approach (Fig 3) and compared the candidate’s model structure with that of the well-characterized hormone receptors DAF-12 in *A. ceylanicum* [14,15] (DAF-12_Ace_) and *S. stercoralis* [15] (DAF-12_Sster_), for which structural and crystallographic information is available. Additionally, we generated a model of *C. elegans* (DAF-12_Cele_) to be used as reference since there is abundant functional information available.

**Fig 3.**
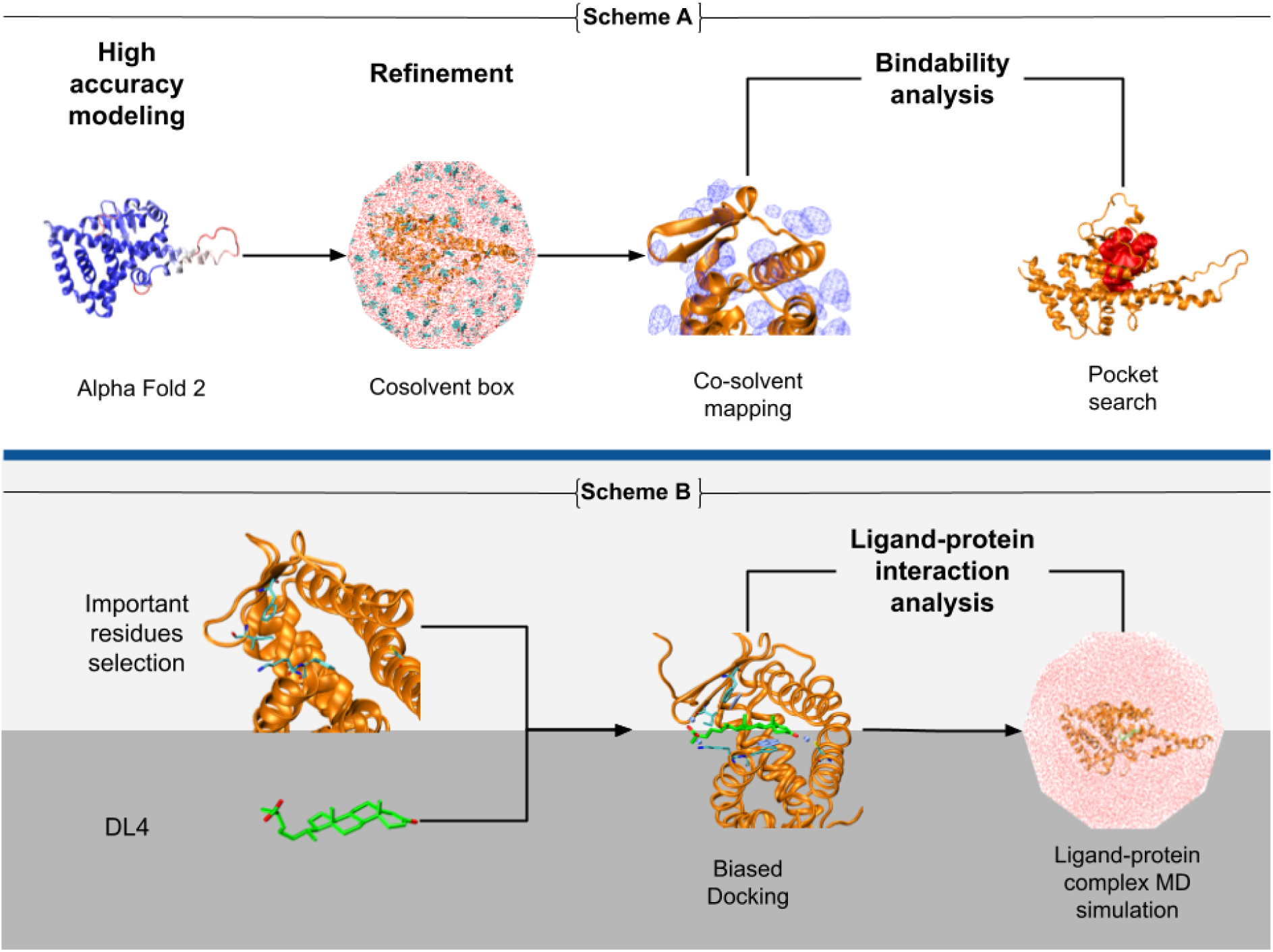
Structural bioinformatics approach employed in this study. Scheme A outlines the procedure for identifying the active site in DAF-12Minc, which involves three-dimensional modeling, refinement using molecular dynamics with mixed solvents, hot sports identification, and cavities characterization. Scheme B demonstrates the approach used to identify significant ligand-protein interactions through guided molecular docking, followed by refinement using molecular dynamics simulations.

Due to the lack of crystal structures for the candidates, we employed AlphaFold2 [16] to model the DAF-12_Minc_ structure from the amino acid sequence of Minc3s00834g17902, which was left available in Target Pathogen Server [17] (S6 File), together with the protein’s annotation and it’s assessment as a potential drug target. To assess the stability of the hormone receptor model, we performed three replicas of 1 microsecond of Molecular Dynamics simulations (MD) and compared it with previously reported structures [14,15]. Our results revealed that this protein had similar secondary structure composition and the fold of the hormone domain remained stable during the simulations (S14 Fig and S2 Table), with slight differences primarily in the size of the H11 helix of DAF-12_Minc_ compared to DAF-12_Sster_ [15], DAF-12_Ace_ and DAF-12_Cele_ (S15 Fig).

Moreover, we found that while all reference structures contain two helices (H8 and H9), DAF-12_Minc_ has only one helix covering the same space as H8 and H9, but with equivalent interactions and folding properties. Subsequently, we used Fpocket [18] to identify pockets that were consistent across the references and models (S2 Table). Interestingly, all identified pockets were formed by a 3-layer α-helical sandwich-like array (S16 Fig).

Taking into account the DA ligands are mainly hydrophobic, we used molecular dynamics with mixed-solvents to obtain a detailed description of the potential protein-ligand interaction hotspot at the DAF12 binding site [19]. First, we estimated the predictive potential of phenol molecules to identify high occupancy regions in DAF-12_Sster_ and compare the key interactions with DAs-like compounds (S17 Fig) [15]. The two highly populated regions in the DAF-12_Sster_ binding site correlates with position of the aliphatic rings of the DA acids, validating the ability of phenol molecules for mapping hydrophobic interactions [19]. In the case of DAF-12_Minc_, similar hotspots were found at the corresponding ligand binding site, indicating that despite the low correlation observed between the relevant residue position in the multiple alignments, the composition and spatial distribution of the amino acids could be sufficient to interact with the ligand.

Finally, to determine the ability of DAF-12_Minc_ to accommodate DA acids at the binding site, we performed biased docking simulations of DA-like molecule DL4. Using the previously identified hotspots as restraints [20], we were able to determine the optimal ligand pose (Fig 4) confirming that the binding mode of DL4 is similar in DAF-12_Minc_ and DAF-12_Sster_.

**Fig 4.**
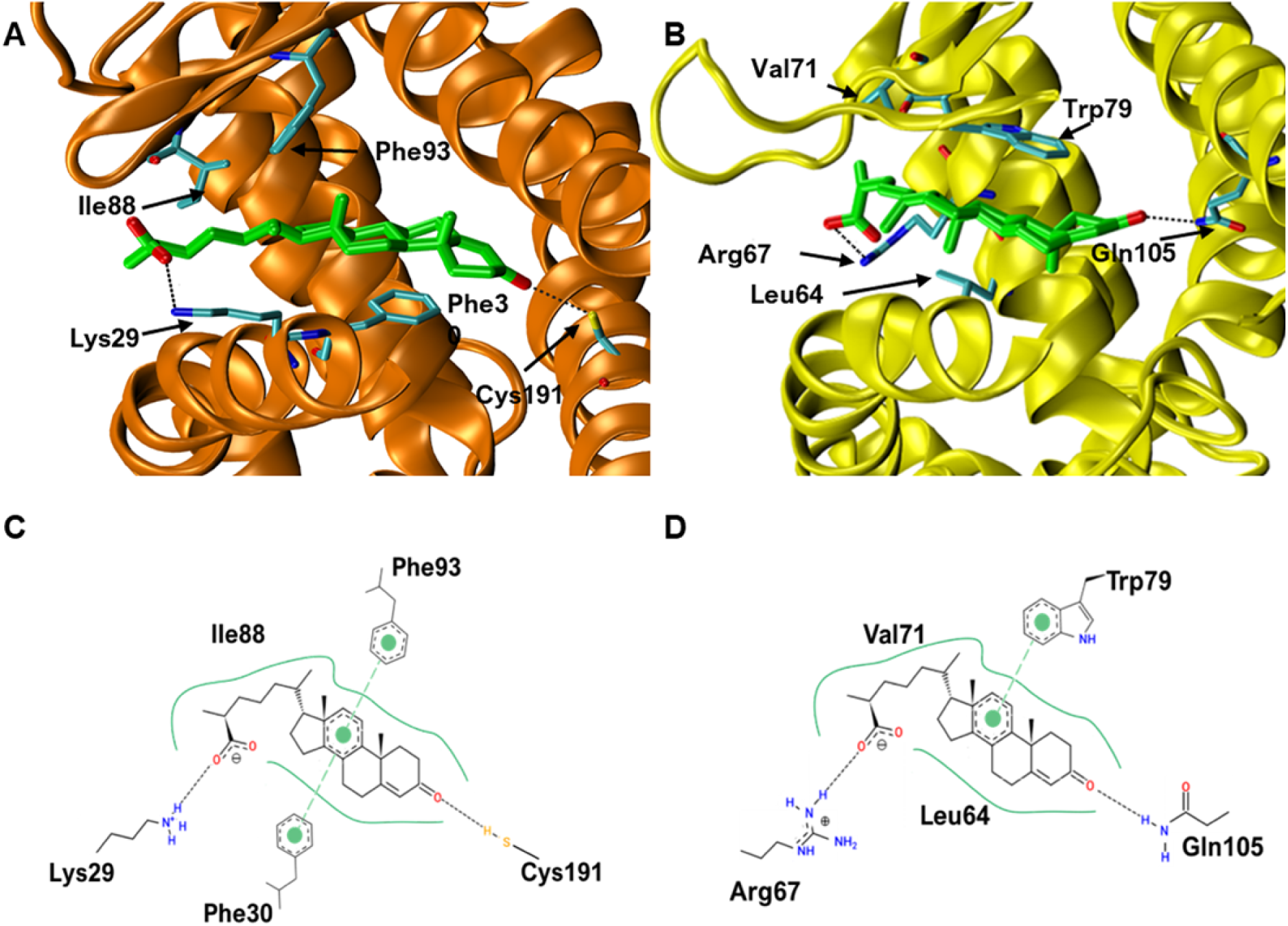
Proposed ligand binding mode for *M. incognita*. This figure illustrates the positioning of the DA ligand at the binding site of hormone receptors in *M. incognita* (Panel A) and *S. stercoralis* (Panel B). The receptor structures are depicted using a cartoon representation, while the ligands and key residues are displayed using a licorice representation. Labels and arrows highlight key amino acids involved in the binding mode. Dotted lines indicate the primary polar interactions between the ligand and the receptor. Panels C and D present 2D diagrams illustrating key interactions between DAF-12Minc-DA and DAF-12Sster-DA, respectively. These diagrams show the aromatic interactions (*π*-stacking), represented by filled green circles, and hydrogen bond interactions, depicted by dotted lines, occurring between key amino acids and the functional groups of DA.

The results of the structural analysis indicated that structural fold, the composition and distribution of residues in the binding site, and the interactions between these residues and ligand atoms were all highly comparable among the orthologous structures. Notably, the mechanism of ligand binding revealed in the DAF-12_Sster_ structure is conserved among DAF-12 proteins.

Based on these findings, we propose that the three-dimensional models of DAF-12_Minc_, DAF-12_Cele_ and DAF-12_Ace_ used in this study are orthologous to the DAF-12_Sster_ protein. This conclusion is supported by the similarities in spatial distribution, number and types of domains, active site characteristics, and ability to bind DA’s-like molecules such as compound DL4. These findings are particularly relevant because DAF-12_Cele_ has been experimentally tested, and the role of DAF-12 has been proved, even though the structure of the receptor was previously unknown. Moreover, the structural similarities further support the efficiency of the employed methodology to identify orthologs.

### *In vitro* validation of the DAF-12 candidate

In order to test if the putative DAF-12_Minc_ gene identified *in silico* codes for a functional DAF-12 nuclear receptor, HEK-293T cells were co-transfected with an expression vector coding the luciferase reporter gene under the control of the atg-7/let-7 promoter of the region CEOP4404 operon containing the atg-7 and let-7 genes (strongly regulated by DAF-12) [6] and an expression vector expressing Minc3s834g17902 or pDAF-12 A1 (codifying DAF-12_Cele_). Cells were then administered with a DA synthetic analogue (Δ^4-24^ DA) at different concentrations. Minc3s834g17902 bound the DA analogue when the concentration was higher than 1µM (Fig 5A).

**Fig 5.**
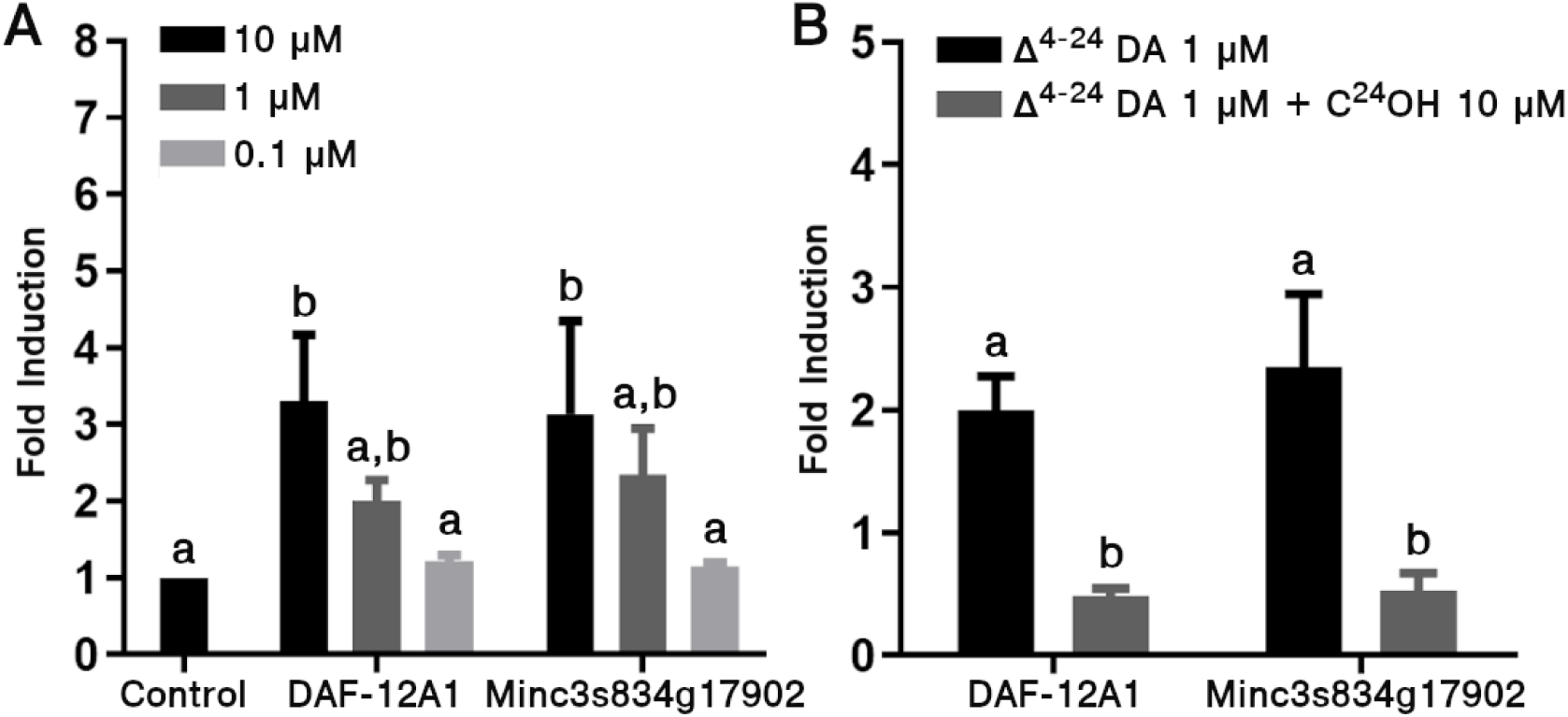
A synthetic analogue of DA and a DAF-12 antagonist regulate Minc3s834g17902 promoter activity. (A) Concentration-response of Minc3s834g17902 and DAF-12A1 (*C. elegans*) with DAF-12 dependent atg-7/let-7-luciferase reporter activity in the presence of increasing concentrations of Δ^4-24^ DA in HEK-293T cells (means ± S.D; n=3); statistical significance was determined by Tukey’s test. In all cases, the same letters are not significantly different (P < 0.05). (B) HEK-293T cells were co-transfected with Minc3s834g17902 and DAF-12A1 with DAF-12 dependent atg-7/let-7-luciferase expression vector. Cells were then treated with 1μM Δ^4-24^ DA in the absence or presence of 10μM antagonist compound. Luciferase fold induction is expressed relative to 1μM Δ^4-24^ DA (100%). Results are means ± S.D (n=3). Statistical significance was determined by Tukey’s test. In all cases, the same letters are not significantly different (P < 0.05).

On the other hand, effects of a DAF-12 antagonist compound were determined against the analogue in a cell-dependent gene expression induction. HEK-293T cells were co-transfected with the same expression vector that was described previously. Cells were then treated with 1μM compound Δ^4-24^ DA in the absence or presence of the antagonist (C^24^OH) [6] at a concentration of 10μM. Fig 5B shows that the antagonist was able to inhibit the effect of Δ^4-24^ DA in the Minc3s834g17902 receptor.

These *in vitro* experiments indicate that the putative DAF-12_Minc_ is capable of promoting gene expression when it binds DA compounds, whereas binding a known antagonist reduces this effect on gene expression. In other words, the putative DAF-12_Minc_ displays the same behaviour as DAF-12_Cele,_ further supporting our strategy to identify orthologous genes.

### *M. incognita* treatment with DAF-12 antagonist compound inhibits egg hatching and increases J2 mortality

It is known that the antagonist alcohol compound C^24^OH treatment induces a delay in *C. elegans* life cycle by binding to the DAF-12 nuclear receptor [6]. In order to determine whether DAF-12_Minc_ affects *M. incognita* development in the same way DAF-12_Cele_ does when treated with this antagonist, egg hatching and J2 mortality were analyzed. Isolated *M. incognita* eggs from galls of infected roots were incubated in the presence of increasing concentrations of antagonist molecule alcohol in 0.2% DMSO or 0.2% DMSO alone as a control for 96 hours. After treatment, barely half of the eggs incubated with 100µM and 200µM had hatched (52 and 46% respectively). The experiment was continued for another extra 24 hours but no significant changes were observed compared to 96 hours results. Significant differences in egg hatching were detected at 96 hours (P < 0,0001) between 100 and 200µM the antagonist molecule alcohol treated ones compared to the control and 50µM the antagonist molecule alcohol treated (Fig 6A).

**Fig 6.**
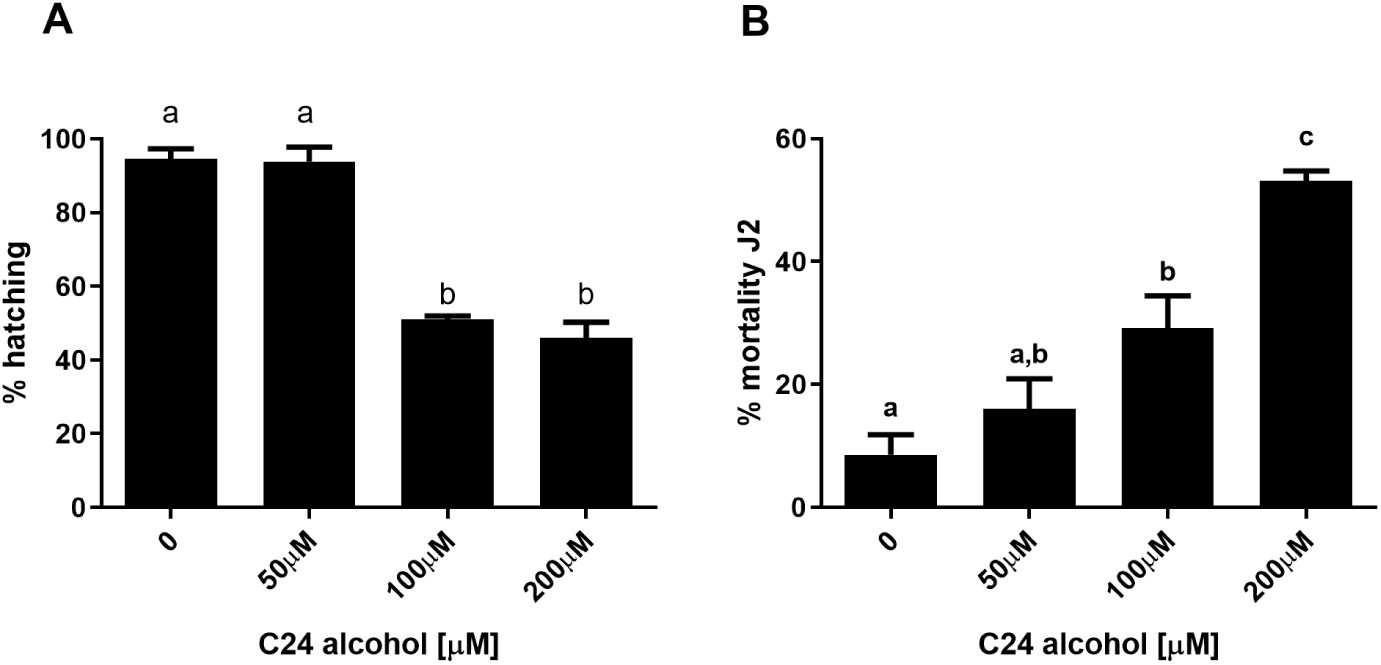
The antagonist compound inhibits *M. incognita* egg hatching and increases J2 mortality. (A) Percentage of hatching was determined after 96 hours of incubation of purified *M. incognita* eggs with increasing concentrations of the antagonist compound C^24^OH (indicated as C24 alcohol) in 0.2% DMSO or 0.2% DMSO. The values shown are the mean of three biologically independent experiments. (B) Percentage of mortality was calculated after 48 hours of incubation of J2 worms in the presence of increasing concentrations of the antagonist compound in 0.2% DMSO or 0.2% DMSO. The values shown are the mean of three biologically independent experiments with 100 eggs. Error bars represent the SEM. Statistical significance was determined using Tukey’s test. In all cases, different letters indicate a significant difference between means (P < 0.05).

To further analyze the effect of the antagonist treatment in *M. incognita* mortality, J2 larvae were incubated in the presence of increasing concentrations of the antagonist molecule alcohol or DMSO as a control and after 48 hours the percentage of dead J2 was calculated. There is a significant dose response effect of C^24^OH in *M. incognita* mortality (Fig 6B). After 48 hours of incubation 16, 28 and 53% of dead nematodes were found in the presence of 50, 100 and 200µM the antagonist molecule alcohol respectively, whereas only 8% of dead J2 were found in the control. All together these results showed that the DAF-12_Cele_ antagonist had a significant effect in the life cycle and mortality of *M. incognita*, thus suggesting the existence of a functional DAF-12 gene in its genome.

### Reduction in migration of the J2 larvae treated with the antagonist compound

To examine the possible role of DAF-12 in the ability of J2 larvae to find new hosts, migration and consequent penetration of the J2 towards the root of tomato plants (a species susceptible to an RKN infection) was analyzed in the presence of the antagonist C^24^OH. Results indicate a reduction of migration and penetration of roots with the addition of the antagonist compound. During the first two hours the percentage of migration to the root did not exceed 40% in the control and 14% in the treated. At the third and fourth hour an increase in migration was observed, reaching up to 75% for control compared to a reduced 30% in the treated (Fig 7A). Moreover, the percentage of J2 that penetrated the root after the third test hour was 25% in the control, higher than the negligible 3% in the treated ones, and after spending an hour more there was no greater difference with respect to these values (Fig 7B). Fig 7C shows the percentage of J2 that migrated after 4 hours of the test, it was observed that 75% of J2 migrated in the control, while those treated with the antagonist barely migrated 30%. Fig 7D shows the percentage of J2 that penetrated the root after 4 hours of assay, and it was found that 25% of J2 entered the control, whereas in those treated with the antagonist, a tiny 3% of worms entered the root. Photos in Fig 7 (panels E-H) show a significant reduction in J2 migration and penetration in the presence of a DAF-12 antagonist compound.

**Fig 7.**
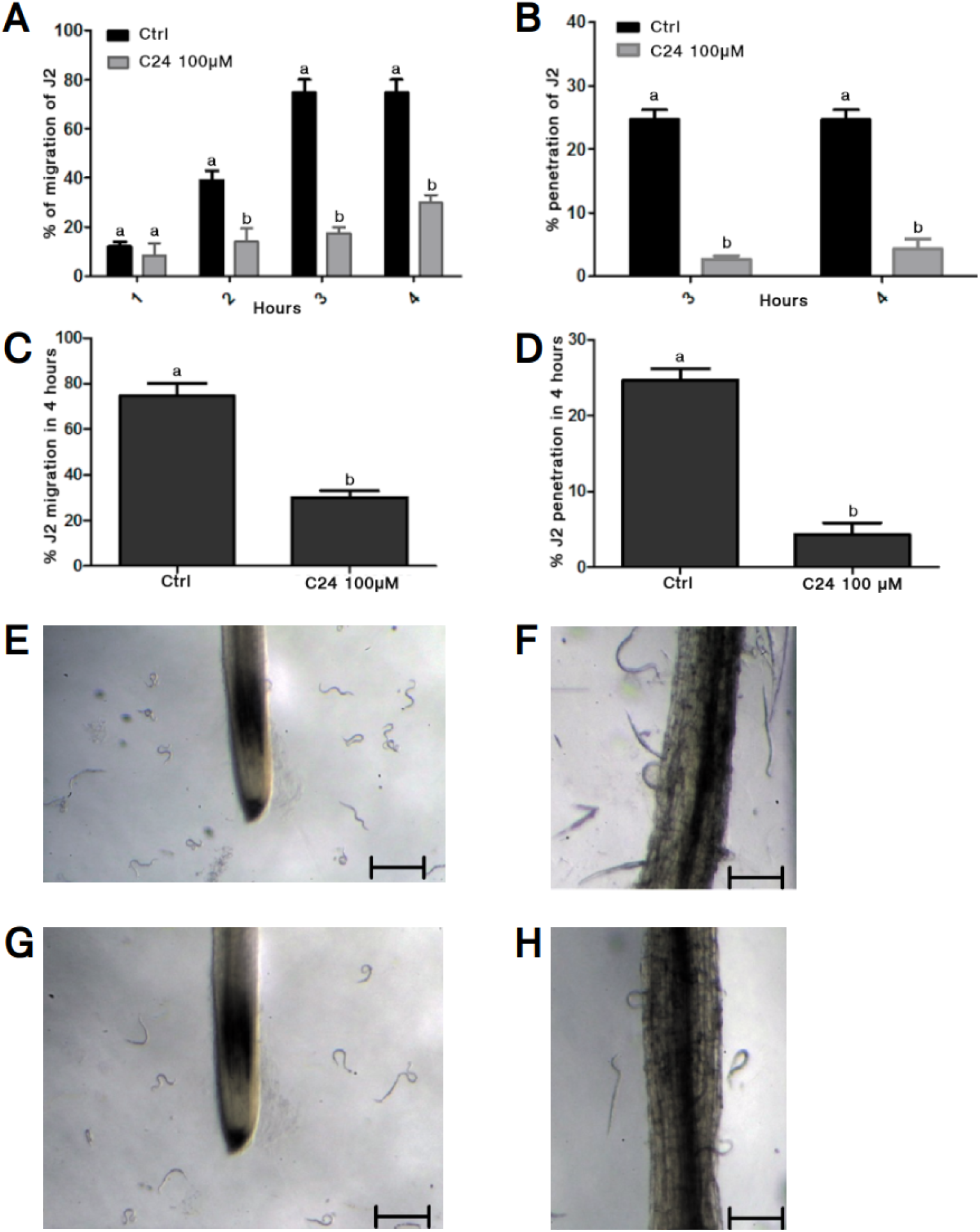
Reduction in migration of the J2s treated with the antagonist C^24^OH. Migration of the J2s treated with the antagonist C^24^OH (C24) in agarose 0,1%. (A) Percentage of migration of J2 towards the root. (B) Percentage of J2 that entered the root. (C) Percentage of J2 that migrated and finally penetrated the root after 4 hours. (D) Percentage of J2 that entered and finally penetrated the root after 4 hours. Photographs displayed migration of the treated J2 in response to tomato root tip at a period of 4 hours (scale bar, 400μm). (E) Photo of untreated tomato plant root with increase 1.5X. (F) Photo of untreated tomato plant root with increase 5X. (G) Photo of tomato plant root treated with 100µM C^24^OH with increase 1.5X. (H) Photo of tomato plant root treated with 100µM C^24^OH with increase 1.5X. Each bar value represents the mean ± SD of triplicate experiments. Statistical significance was determined using Tukey’s test. In all cases, different letters indicate a significant difference between means (P < 0.05).

In summary, this experiment is an indicator that suppressing the activity of DAF-12 affects the ability of J2 larvae to migrate and penetrate plants, which in turn should impede their normal maturation by infecting the plant.

### DAF-12 antagonist interferes with *M. incognita* tomato plant infection

In order to determine whether interfering with DAF-12 not only affects migration and penetration of the plant, *M. incognita* infection was evaluated in tomato plants with or without the antagonist compound. 6 days old tomato plant roots were treated for 48 hours with increasing concentrations of the antagonist molecule alcohol in DMSO 0.2% or DMSO 0.2% as control and then infected with 100 J2. The number of galls and egg masses per plant were determined 35 days after infection. Treatment with the antagonist decreased significantly the number of galls per root system in a dose-response manner. As shown in Fig 8A, at 100µM the antagonist compound produced a 74% reduction in the number of galls. At 200µM the antagonist did not cause an increasing reduction compared to the 100µM treatment (68% reduction in the number of galls). However, treatment with the antagonist at 50µM did not present any significant difference compared to control plants (although there was a tendency of a 25% reduction in the number of galls). Fig 8B shows a 54% and 46% decrease in egg masses per root system when tomato plants were treated with the antagonist compound at 100 and 200µM. No differences in egg masses per root system were found between treatment with antagonist C^24^OH 50µM and control plants. Altogether, these results suggest that *M. incognita* infection is reduced when plants are treated with a DAF-12 antagonist.

**Fig 8.**
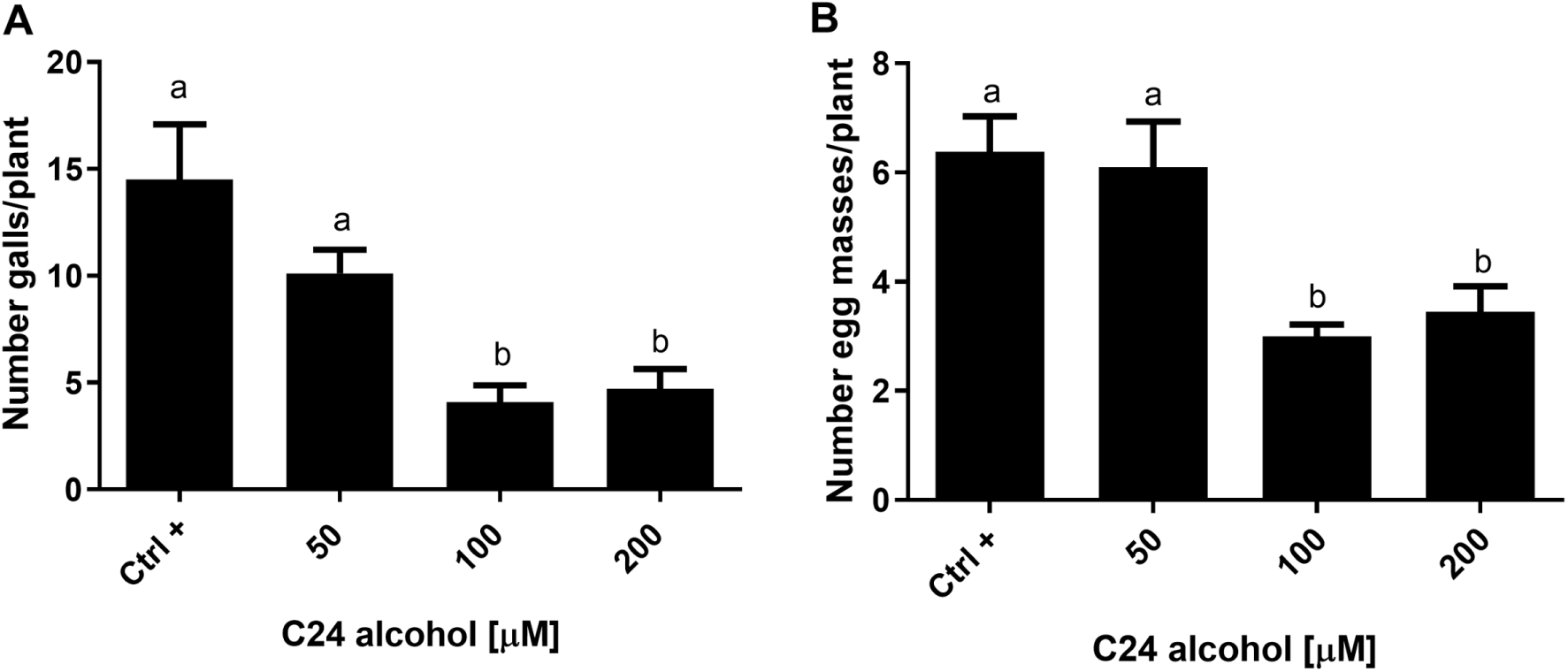
The antagonist compound treatment of tomato plants inhibits *M. incognita* infection. Untreated or treated tomato plants with increasing C^24^OH concentrations (C24 alcohol) were infected with 100 *M. incognita* J2. (A) Number of galls per plant. (B) Number of egg masses per plant. The values shown are the mean of three biological independent experiments with 10 plants. Error bars represent standard deviations. Statistical significance was determined using Tukey’s test. In all cases, different letters indicate a significant difference between means (P < 0.05).

### Tomato plant growth parameters improve under treatment with DAF-12 antagonist compound

Interference with normal DAF-12 activity and the consequential negative effects on *M. incognita* life-cycle and plant infection could imply that plants infected by J2 larvae with malfunctioning DAF-12 should display an overall better plant health condition. To test this, tomato plants treated with 100µM of the antagonist compound C^24^OH in DMSO 0.2% or DMSO 0.2% (Ctrl+) were infected with 100 J2. Uninfected plants were also included (Ctrl-).

Several growth parameters were determined, including fresh weight, root and stem size and number of leaves 35 days after infection. Fig 9 shows that *M. incognita* infected plants pretreated for 48 hours with the antagonist show a statistically significant growth according to the observed growth parameters, similar to the uninfected ones, when compared to the untreated infected plants (Ctrl+). Specifically, root size in treated plants was 17% lower than uninfected plants, while average root size in untreated infected plants was 61% lower than the uninfected control (Fig 9A). No statistically significant differences were observed in stem size between treated and untreated infected plants (Fig 9B). On the other hand, there was a statistically significant 50% increase in leaves number and 75% in fresh weight between treated and uninfected plants (Fig 9C and Fig 9D). Moreover, no statistically significant differences in all analyzed parameters were found between the 100µM C^24^OH treated infected plants and the uninfected ones. In addition, when uninfected plants were treated with the antagonist compound no significant differences were found compared to the uninfected and untreated ones (data not shown). Altogether, these results could be indicative that the health improvement observed in treated plants infected with J2 was due to the interference with DAF-12 role in development by the antagonist compound on *M. incognita*.

**Fig 9.**
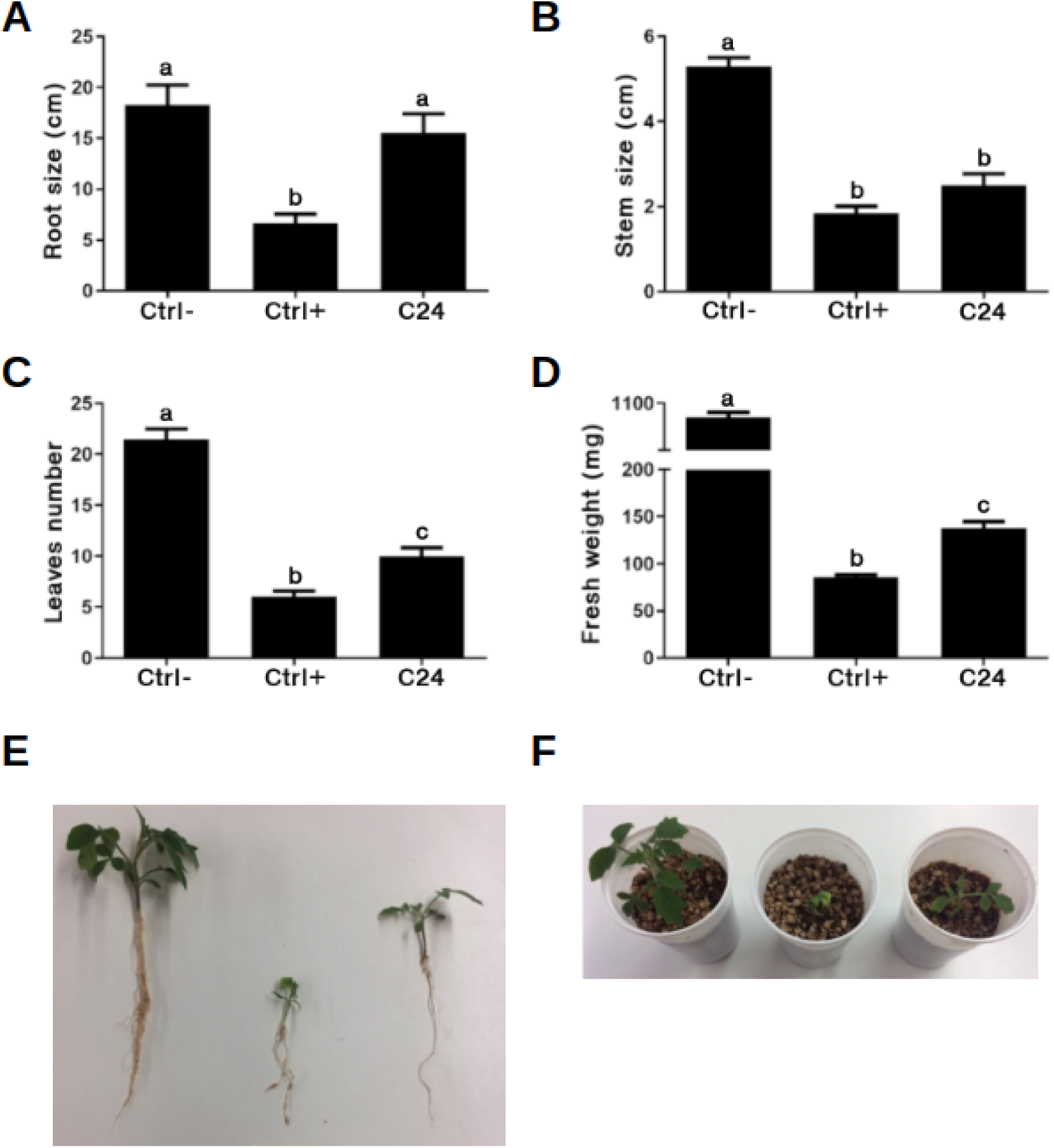
Improvement in plant growth parameters in C^24^OH pretreated tomato plants infected with *M. incognita*. Different plant growth parameters were determined in uninfected (Ctrl-) and treated (C24) or untreated (Ctrl+) with 100µM C^24^OH infected tomato plants. (A) Root size. (B) Steam size. (C) Number of leaves. (D) Fresh weight. (E) Representative uninfected and treated or untreated with the antagonist at 100µM infected tomato plants. (F) Plants shown in the previous panel in their pots. The values shown in the bar plots are the mean of three biologically independent experiments with 10 plants. Error bars represent SEM. Statistical significance was determined using Tukey’s test. In all cases, different letters indicate a significant difference between means (P < 0.05).

### Embryonic lethality in *M. incognita* induced by DAF-12 antagonist compound

In order to study the long term effect of the antagonist compound in the M. incognita life-cycle we analyzed the hatching capacity of eggs recovered from tomato plants treated with the compound. To this end, eggs developed in galls of plants treated with 100µM C^24^OH or DMSO were collected and incubated in water for 8 days at 28°C and the percentages of J2 hatching were determined. As shown in Fig 10, 54% of the eggs collected from the antagonist compound treated tomato plants were unable to hatch in standardized lab conditions. This result supports the idea that inhibition of DAF-12 by the antagonist has an effect not only in the infection by *M. incognita*, but also a long term effect in the inhibition of offspring hatching. Two possible scenarios could explain this result: either the antagonist compound affects the development of the female and in consequence its eggs viability, or the antagonist is still persistent after 45 days of plant treatment and is able to affect embryos during their development in the egg sacks.

**Fig 10.**
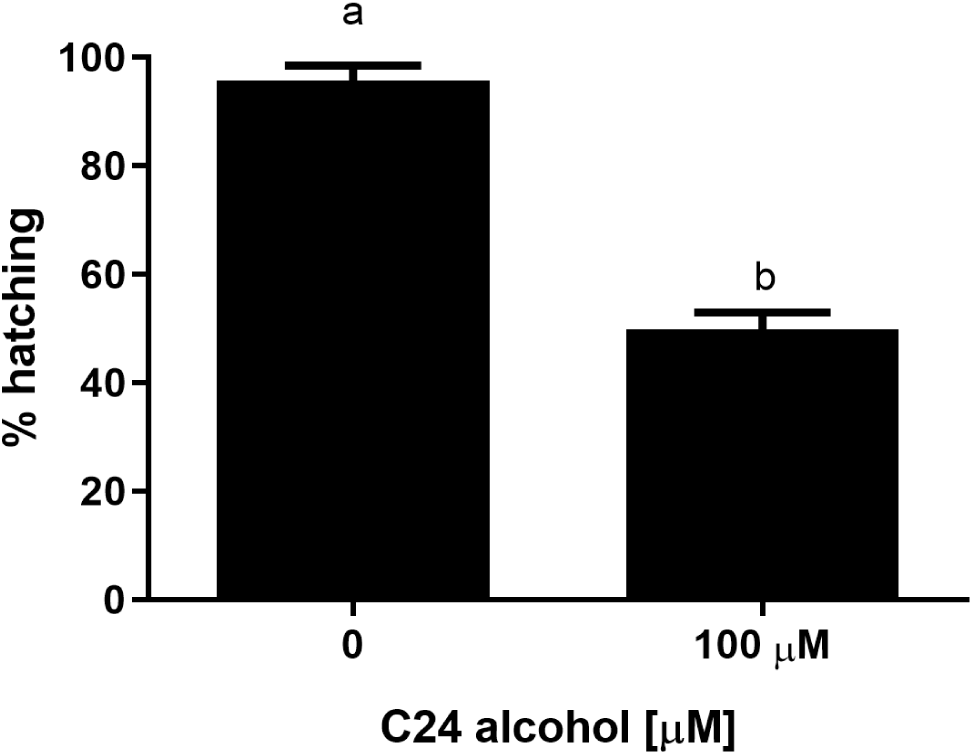
The antagonist compound reduces *M. incognita* hatching of eggs purified from the antagonist-treated plants. Percentage of hatching of eggs purified from galls of untreated and 100µM C^24^OH (C24 alcohol) treated plants after 8 days of water incubation. Values shown are the mean of three biologically independent experiments with 100 eggs. Error bars represent standard deviations. Statistic significance was determined with t-test. Different letters indicate a significant difference between the means (P < 0.05).

## Discussion

DAF-12 is a type of nuclear receptor that responds to hormone signals in nematodes and plays a critical role in regulating the maturation of larvae. While this protein was initially identified in the free-living species *C. elegans*, it has also been found in parasitic worms such as *S. stercoralis*, where it performs a similar function in regulating the progression towards adulthood. Given the vital role of DAF-12 in nematode biology, it has emerged as a promising molecular target for controlling the spread of plant-parasitic nematodes like *M. incognita*, which can cause significant damage to crops. However, despite previous attempts, no DAF-12 ortholog has been found in this parasite to this day.

The identification of gene orthologs is one of the biggest challenges in bioinformatics. Although tools based on local alignments like Blast are efficient to find orthologous genes in most cases, when the organisms are too divergent results may be hard to interpret and may lead to erroneous conclusions. Therefore, an approach that takes in consideration different aspects of a protein such as sequence conservation, domain presence, phylogenetic relationships, structure, function or ligand-binding capacity will return more reliable results. In the present study, we developed a new methodology to identify orthologs with the intention of finding DAF-12 in *M. incognita* that allowed us to address this problem.

An initial HMMER search on the *M. incognita* proteome using the ligand-binding domain and DNA-binding domain allowed us to reduce the number of candidates from more than 40 thousand protein sequences. This initial filter has a double advantage: it significantly reduces the number of sequences to a reasonable number (52 sequences) and it ensures that the remaining ones contain all the necessary domains to carry on the DAF-12 function, a requirement to be the ortholog. However, this begs the next question: which of the 52 sequences are orthologs and which ones are just paralogs with the same domains? By generating a phylogenetic tree with all 52 candidates and validated DAF-12 sequences from other nematodes, we can be certain that those *M. incognita* sequences that have a closer phylogenetic relationship to the reference sequences are most probably DAF-12 orthologs.

The advantage of phylogenetics in contrast to sequence similarity is that the first one can group together sequences that are very divergent, if they still share some key residues that are clearly conserved. This approach led us to narrow down the number of DAF-12_Minc_ candidates to 3, given the consistent result of having the same 3 sequences grouped with the DAF-12 reference sequences both in the protein and RNA phylogenetic trees. This consistency in the phylogenetic results further supports the final 3 candidates obtained.

Once the phylogenetic analysis was performed, what remained to be determined was whether these proteins fold like DAF-12_Cele_, DAF-12_Ace_ and DAF-12_Sster_, and if they were capable of binding the corresponding ligand: dafachronic acids. Our research led us to observe the structural similarity between proteins once we modeled and aligned them. After a molecular dynamics refinement, the pocket prediction showed the best scored pocket in the same place as the protein crystals used as reference. Moreover, in order to obtain as much information as we could we proceeded to map the possible active site using co-solvent sites data retrieved from MD simulations. This methodology has demonstrated high predictive capabilities before [21] [22], and in the present work it contributed to identifying the spatial distribution of sites where important interactions occur which are very similar among the proteins. Those interactions are mediated by amino acids with similar physicochemical characteristics. One of the most important ones is the π-stacking interaction carried out by a Trp in DAF-12_Sster_ [15], a residue that was not in the same place at the DAF-12_Minc_ binding site but that appears to be replaced by a Phe right on top of the DAF-12_Minc_ pocket. This residue seems to be important because the molecular dynamics and co-solvent mapping results showed that it moves until it is stabilized by a phenol methyl group found perpendicularly to the aromatic residues (Fig 4). Also, as mentioned before, in the mutated DAF-12_Sster_ Trp absence leads to no biological activity. This could be due to π-stack interactions that stabilize the ligand into the binding site [15].

Once we were able to determine the structural similarities and binding capabilities between the receptors, we performed biased docking analyses using the binding hotspots as restraints. These analyses yielded the best binding position for the canonical ligand and even when the orientation of the DA appeared to be upside down, the binding mode and energy among the results showed that in the DAF-12_Minc_ system, DA-like molecules were capable of forming bonds with the selected residues. Moreover, the interaction analysis led us to conclude that all four proteins were capable of accomplishing the same kind of interaction with the ligand, even when there were substantial differences between the amino acids in the active site and the positions of the ligand.

To this point, it was possible for us to go from an entirely predicted proteome of *M. incognita* to a modeled protein structure employing only bioinformatic tools. To be completely sure that our candidate corresponds to a functional DAF-12 ortholog we performed a luciferase assay to determine whether the DAF-12_Minc_ candidate was capable of promoting gene expression when it binds its ligand. This experiment further supports the hypothesis of the orthologous DAF-12 candidate, since it demonstrated that the protein binds to the predicted promoter in the presence of dafachronic acids.

The final step in assessing whether our candidate is a functional DAF-12 gene, *in vivo* experiments were performed to test if the gene plays a significant role in cell-cycle, larval development and host infection (in the case of parasitic worms), as it does in other nematode species. Suppression of normal protein activity via a DAF-12 antagonist compound had a significant effect both in the parasite and the host plant. In the parasite, suppression of DAF-12 activity resulted in an increase in larval mortality and embryonic lethality; and a decrease in egg-hatching, larval migration, root penetration and plant infection. In the host plant it resulted in an overall improvement in the plant’s health, which was measured by a number of parameters such as root size, stem size, number of leaves and fresh weight. Altogether, these experimental results point to the fact that our DAF-12 candidate is key to life-cycle and plant infection, as it was expected.

In summary, an initial HMM-based search combined with phylogenetic analysis allowed us to narrow down the number of possible DAF-12 orthologs to 3. Structural analysis indicated that these candidates have a very similar structure to other DAF-12 proteins described in other nematodes, and that they are capable of binding their expected ligands, the dafachronic acids. *In vitro* experiments suggested that these candidates can promote gene expression as a response to their ligand. *In vivo* experiments showed that these genes have the same functionality (larval development and host infection) that DAF-12 has in other nematodes. In other words, we were able to identify an orthologous DAF-12 gene in *M. incognita*, which demonstrates that our methodology based on bioinformatic tools is a powerful, effective and efficient way to find orthologous genes among the sequences.

## Materials and methods

### Computational methods and i*n silico* analysis

#### Data mining and phylogenetic analysis

The proteome and CDS transcripts of *M. incognita* were retrieved from BioProject ID PRJEB8714 [11] from the online WormBase database [23]. In order to filter the sequences a data mining on the *M. Incognita* proteome with HMMER (http://hmmer.org/) was performed, using the *zf-C4* (PF00105) and *Hormone_recep* (PF00104) Pfam HMMs for DAF-12 and with p450 (PF00067) for DAF-9 [24]. Nucleotide sequences corresponding to the filtered protein sequences were extracted from the CDS transcriptome. Nucleotide and protein sequences of both genes were aligned with MAFFT [25], the alignment was manually curated with Jalview [26] and finally trimmed with BMGE [27]. Then, phylogenetic analyses were performed using the maximum likelihood method by PhyML software [28] using the Booster bootstrap branch support [29]. Substitution models for nucleotide and amino acid sequences were automatically chosen with SMS [30]. Phylogenetic trees were generated and visualized with the R package ggtree [31]. In addition, the gene and protein sequences from other nematode species (as well as the outgroup sequences) used in these analyses were retrieved from UniprotKb [32]. Pairwise identity matrices from nucleotide and amino acid sequences as well as heatmaps were generated with Python3 libraries matplotlib and seaborn.

#### Protein and ligand structures

The structures of DAF-12_Sster_ (PDBid 3GYT [15]) and DAF-12_Ace_ (PDBid 3UP0 [14]) were obtained from the RCSB Protein Data Bank (http://www.rscb.org/) and were used as benchmarks for this study. Dafachronic acid delta 4 (DL4) was extracted from the 3GYT file. All sequences were subjected to homology modeling using AlphaFold2 [16].

#### Molecular dynamic (MD) simulations

Each system was first optimized using a conjugate gradient algorithm for 5000 steps, followed by two thermalizations. The first one a 10 ps long constant volume MD at 10 K and a restraint of 50 kcal/mol; and the second one in which the first 100 ps were used to gradually raise the temperature of the system from 10 to 300 K (integration step = 0.0005 ps/step). The heating was followed by a 250 ps long constant temperature and constant pressure MD simulation to equilibrate the system density (integration step = 0.001 ps/step). Next, a second equilibration MD of 500 ps. was performed, in which the integration step was increased to 2 fs and the force constant for restrained alpha-carbons was decreased to 2 kcal/mol/Å. Finally, a 10 ns long MD simulation was carried out with no constraints and the “Hydrogen Mass Repartition” technique [33], which allows an integration step of 4 fs, and these conditions were kept for all the subsequent production of 1µs long MD runs with 10 steps per ns having 10000 steps per simulation. All simulations were performed with the Amber package using the ff19SB force field [34] for protein residues and OPC.

For mixed solvents MDs 1M phenol in a water box was used as a cosolvent, while for protein-ligand complex MDs water molecules were used as solvent.

Pressure and temperature were kept constant using the Monte-Carlo barostat and Langevin thermostat respectively, using the default coupling parameters. All simulations were performed with a 10 Å cut-off for non-bonded interactions, and periodic boundary conditions using the Particle Mesh Ewald summation method for long-range electro-static interactions. Visualizations and image rendering were performed with VMD software [35].

### Ligand binding site identification and characterization

We used the Fpocket [18] program to determine the features of pockets in the reference structures before (raw AlphaFold2 models) and after (refined AlphaFold2 models) Molecular Dynamics simulations.

Cosolvent clusters were calculated using an in-house developed TCL script that takes as input the trajectory results of each mixed solvents MD. For cluster building parameters were set at a step of 1 (10000 frames in total), cluster radius of 1.4 A (meaning all phenols inside the radius threshold were taken as part of each cluster), phenol dummy atoms (DU) mass center for aromatic interactions and oxygen atoms (O1) as hydroxide interactions. These interactions were retrieved by keeping those sites where phenoles stayed for at least the 10 % of the dynamic simulation time.

### Molecular docking simulations

Protein and ligand structures were prepared in the PDBQT file format. The grid size and position were chosen so that they include the whole amino acids from the binding site. This was achieved by placing the grid center in the geometric center of the ligand, and extending its size 20 Å in each direction, and using a grid spacing of 0.375 Å. To calculate poses and energies among the runs AutoDock4 [36] was used where the genetic algorithm parameters for each conformational search run were kept at their default values. For each calculation, always 100 different docking runs were performed, and the resulting 100 poses were clustered according to the pyranose ring heavy atom RMSD using a cut-off of 1.5 Å with the quality threshold algorithm implemented in VMD software.

### Reporter gene assay

HEK-293T (human embryonic kidney) cells were cultured in DMEM (Dulbecco-modified Eagle’s medium) supplemented with 10% (v/v) FBS (fetal bovine serum) containing 100 IU/ml of penicillin and 100 μg/ml of streptomycin at 37 °C in a humidified atmosphere with 5% CO2 and 95% O2. Reporter gene assays were performed by transfecting HEK-293T cells with polyethylenimine (PEI) in 12-well plates with serum-free DMEM as described in Dansey *et al.* 2015 [6]. On the one hand, the cells were transfected with the luciferase reporter gene under the control of the atg-7/let-7 promoter in the CEOP4404 operon region that contains the tightly regulated atg-7 and let-70 (Let/luc) genes by DAF-12 and an expression vector pCMVmycDAF-12, together with 450ng of Minc3s834g17902 or DAF-12A1, 1.25ng Let/luc and 219ng of β-galactosidase.

5 hours after the transfections, the cells were washed and the culture medium was replaced by serum-free medium, and the ligands (applied from a stock solution in DMSO) were added. 16-20 hours after the addition of ligands or vehicle control, luciferase and β-galactosidase activities were measured. Each experiment was repeated three times and the induction times were relativized to the unstimulated control.

### Meloidogyne incognita experiments

#### Egg hatching and J2 development

*M. incognita* eggs were collected from infected summer squash (*Cucurbita maxima*) plants growing in greenhouses located at the experimental field of the Faculty of Agronomy, Universidad Nacional del Litoral, Esperanza city, Province of Santa Fe, Argentina (latitude 31° 26’ 31’’ S and longitude 60° 56’ 27’’ O) [37]. Light microscopic examinations of the perineal pattern of female root-knot nematodes were made according to Hartman and Sasser procedure and confirmed the exclusive presence of *M. incognita* [38]. To collect *M. incognita* eggs from roots, Hussay and Barker method was used [39]. For egg hatching experiments, 100 eggs were incubated in presence of increasing concentrations of the antagonist molecule alcohol (50, 100 and 200µM) in 0.2% DMSO or 0.2% DMSO alone as control in a P24 plate at 28 °C for 96 hours. The percentage of hatching was calculated as J2/(J2+egg) x 100, experiments were performed in triplicate. Second stage juveniles (J2) were obtained by incubation of egg masses in a hatching chamber set at 28 °C for 48 hours [40].

#### J2 mortality assay

100 J2 were incubated in the presence of increasing concentrations of the antagonist molecule alcohol (50, 100 and 200µM) in 0.2% DMSO or 0.2% DMSO alone in a P96 plate at 28 °C for 48 hours in a final volume of 250µL. The mortality of J2 was determined by adding 5μL of NaOH 1N to the solution, and J2 that changed shape from straight to curled or hook-shaped within 1 minute were considered viable and if they were motionless they were considered dead [41]. Each treatment was done in triplicates and values are the mean of two different biological replicates.

#### Migration and penetration assay with J2 treaties

Five individual sterile 5-day-old tomato root tips of 1.5cm in length with few root hairs were placed onto a glass with 60μL of 0.1% of agarose containing 40 J2 that had been previously treated for 1 hour with the antagonist molecule alcohol in 0.02 % DMSO or 0.02 % DMSO alone as a control at 2mm from the root tip at 20 °C. J2 were allowed to migrate or penetrate the root for 4 hours and photographed every hour to determine the percentage of worms found at distance less than 0.5mm from the root. In this experiment, images were taken with a ZTX-E Numak stereoscopic binocular at 10X magnification.

### Tomato plant experiments

#### Plant growth, treatment with the antagonist molecule alcohol and nematode infection

Tomato seeds were surface sterilized with 0.1% NaClO for one minute, rinsed 3 times with sterile distilled water, seeded in Petri dishes containing 8% agar and germinated in a greenhouse chamber at 28 °C for 7 days with a 16 hours light and 8 hours dark cycle. Tomato seedlings were transferred singly to 3mm filter paper pouches, placed in a 10 x 20 cm plastic bag and arranged in a rack in a vertical position [42]. 6 days old tomato roots were treated with 300µL of different concentrations of the antagonist molecule in 0.2% DMSO or 0.2% DMSO alone using a micropipette 48 hours after transfer and infected with 100 J2 or with 100µL of water as a control 24 hours later. Seedlings were maintained in a greenhouse chamber at 28 °C in a 16 hours light and 8 hours dark cycle for 4 more days and then transferred to pots containing an autoclaved (80:20) mix of dune sand and vermiculite. Plants were watered twice a week with 5mL of Hoagland solution [43] and tap running water alternatively. Experiments were done in triplicates with 10 plants per treatment.

#### Plant post infection analysis

35 days post infection the number of galls per plant was calculated. Also, tomato plant roots were stained with phloxine B (0.15 g L^-1^) for 30 minutes to estimate the number of egg masses per plant. In order to assess the overall plant health, different plant growth parameters such as root and stem size, plant fresh weight and leaves number were analyzed 35 days post infection.

#### Nematode post infection analysis

100 eggs obtained from infected tomato plants treated with 100µM the antagonist molecule alcohol in 0.2% DMSO or 0.2% DMSO alone were placed in a P24 plate at 28 °C for 8 days. The percentage of hatching was calculated as J2/ (J2+egg) x 100, experiments were performed in triplicate.

### Statistical analysis

Data were analyzed by using one-way ANOVA and Tukey’s multiple comparison test functions and two-tailed t-test for pairwise comparisons in GraphPad Prism 6 software. p-values below 0.05 were considered statistically significant.

## Supporting information

Fastas and excel files

## Acknowledgements

We thank UBA-FCEN-QB-Cluster for granting use of computational resources which allowed us to perform most of the computational analyses included in this work.

