## Supplementary material for "Hidden Markov Models based search in combination with structural bioinformatics pipeline leads to the identification of DAF-12 distant orthologous in *Meloidogyne incognita*": Fastas and excel files

### Supplementary Materia

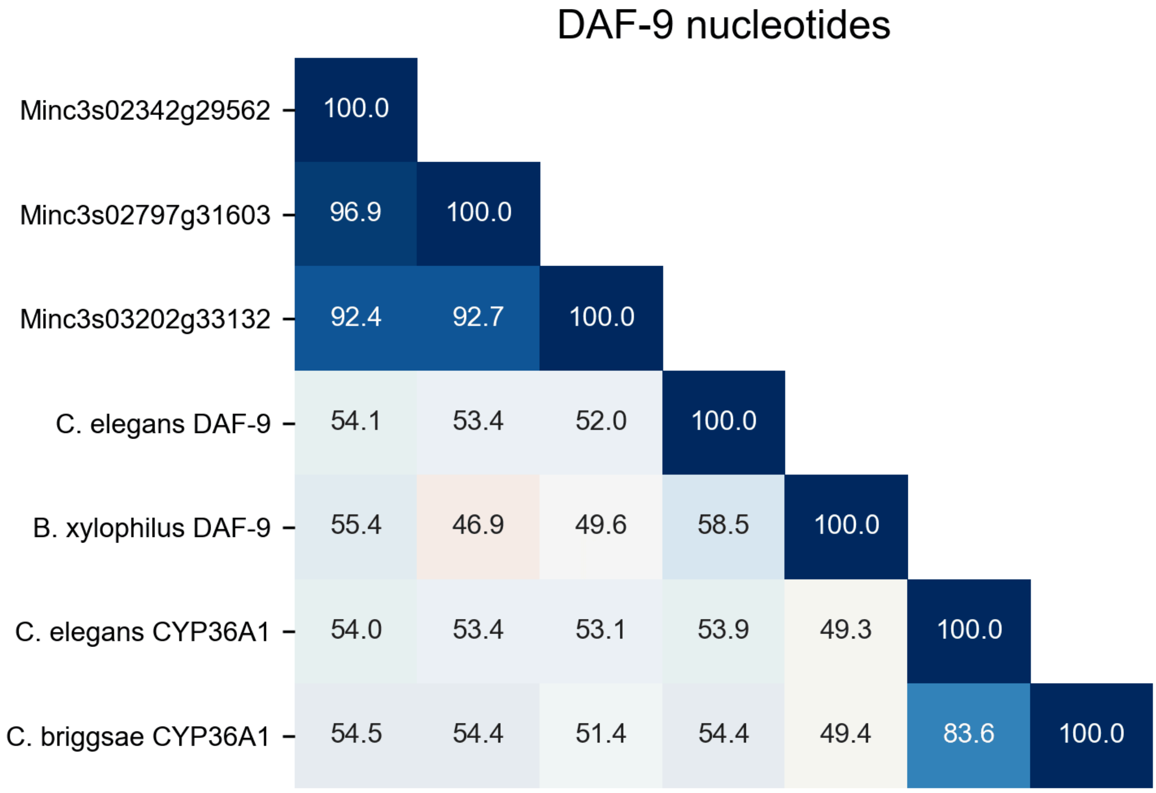


**S1 Fig. DAF-9 nucleotide identity matrix.** Numbers indicate the percentage of pairwise identity between sequences. Colour scale goes from lower sequence identity (red) to higher sequence identity (blue).

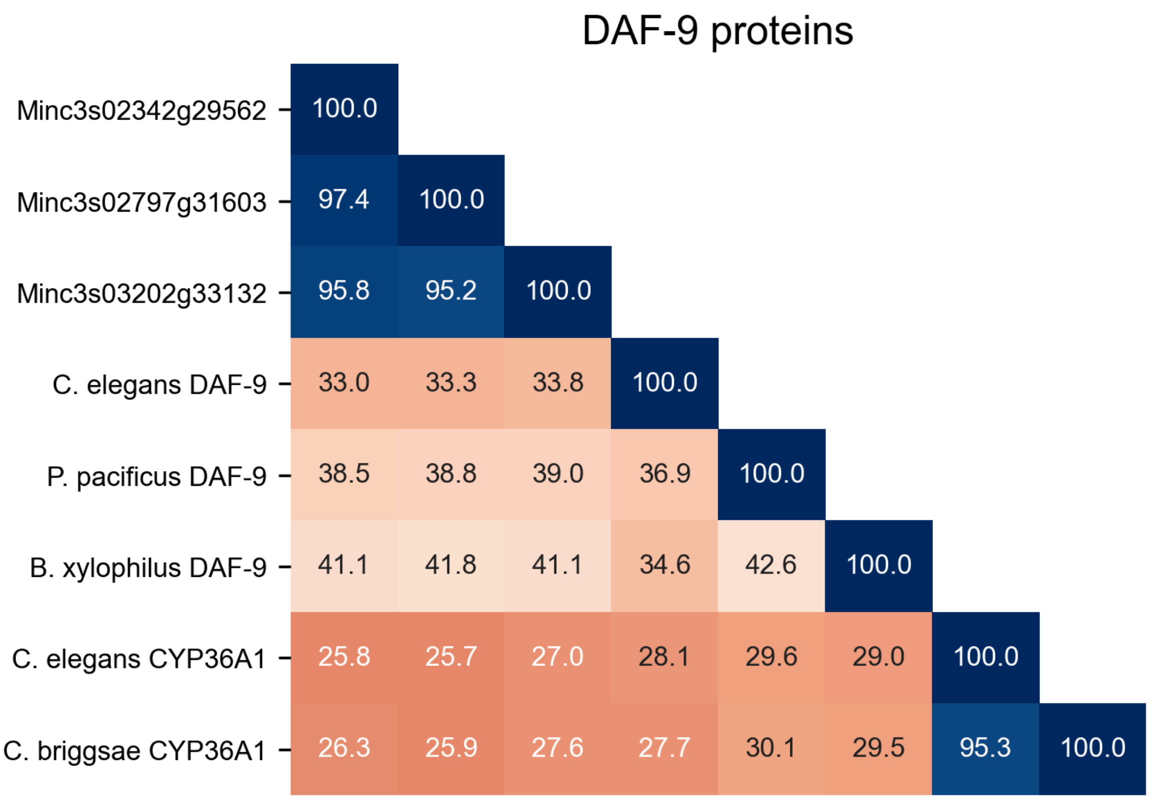


**S2 Fig. DAF-9 protein identity matrix.** Numbers indicate the percentage of pairwise identity between sequences. Colour scale goes from lower sequence identity (red) to higher sequence identity (blue).

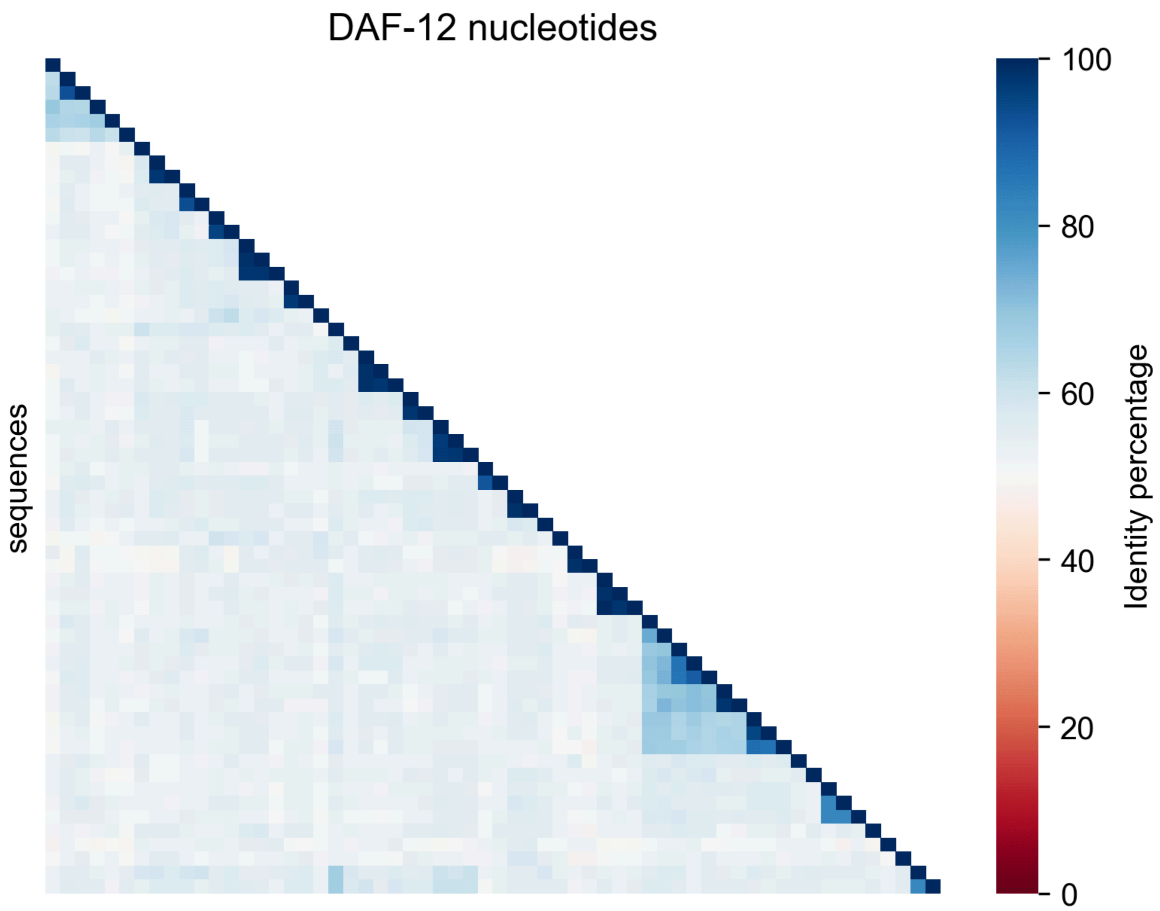


**S3 Fig. DAF-12-*like* nucleotide sequences heatmap.** Colour scale goes from lower sequence identity (red) to higher sequence identity (blue).

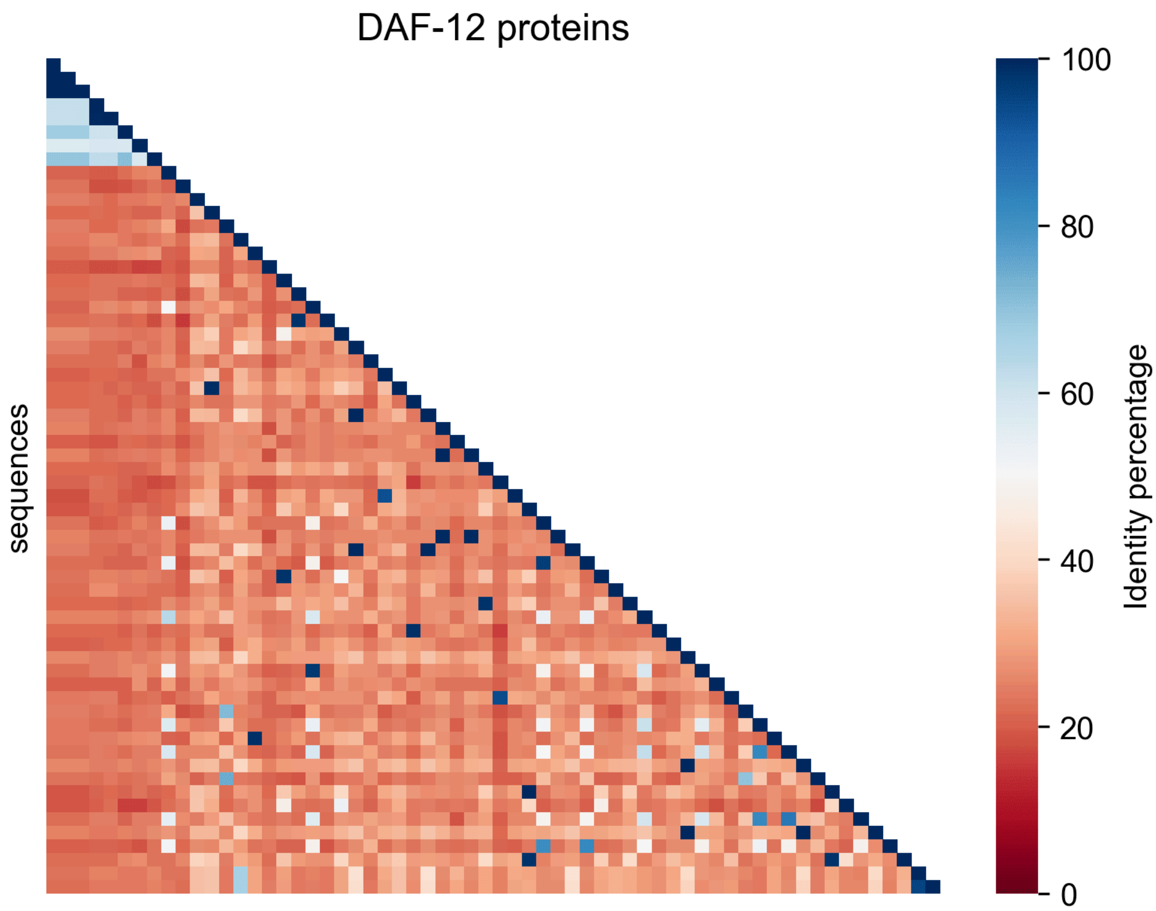


**S4 Fig. DAF-12-*like* protein sequences heatmap.** Colour scale goes from lower sequence identity (red) to higher sequence identity (blue).

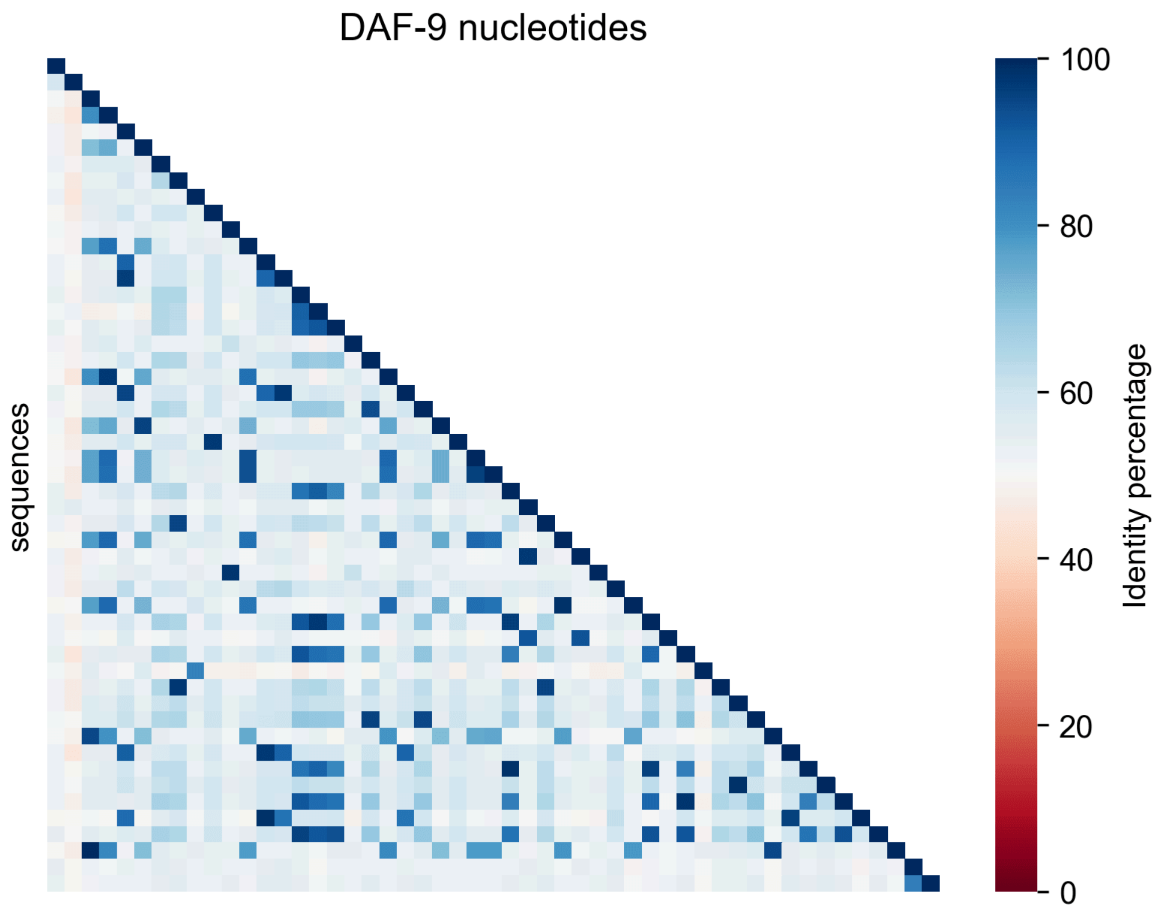


**S5 Fig. DAF-9-*like* nucleotide sequences heatmap.** Colour scale goes from lower sequence identity (red) to higher sequence identity (blue).

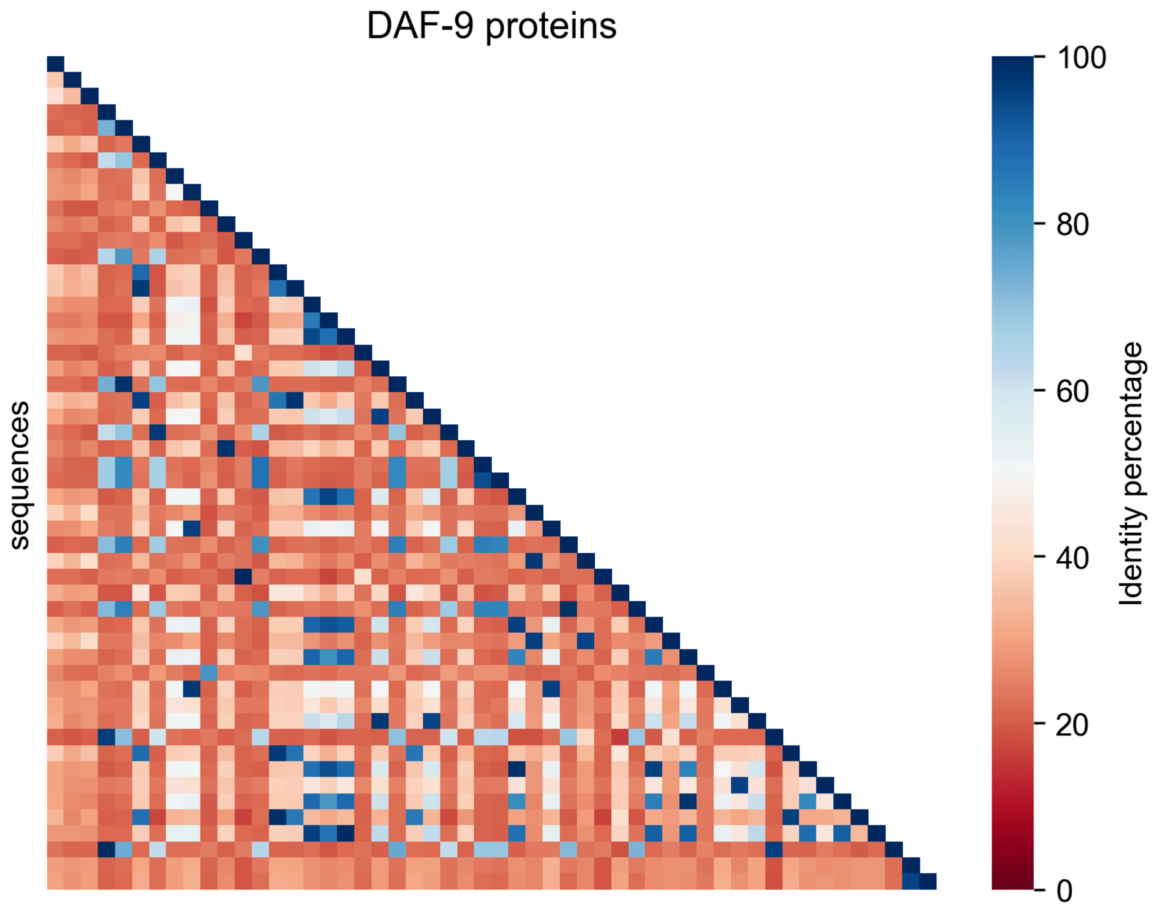


**S6 Fig. DAF-9-*like* protein sequences heatmap.** Colour scale goes from lower sequence identity (red) to higher sequence identity (blue).

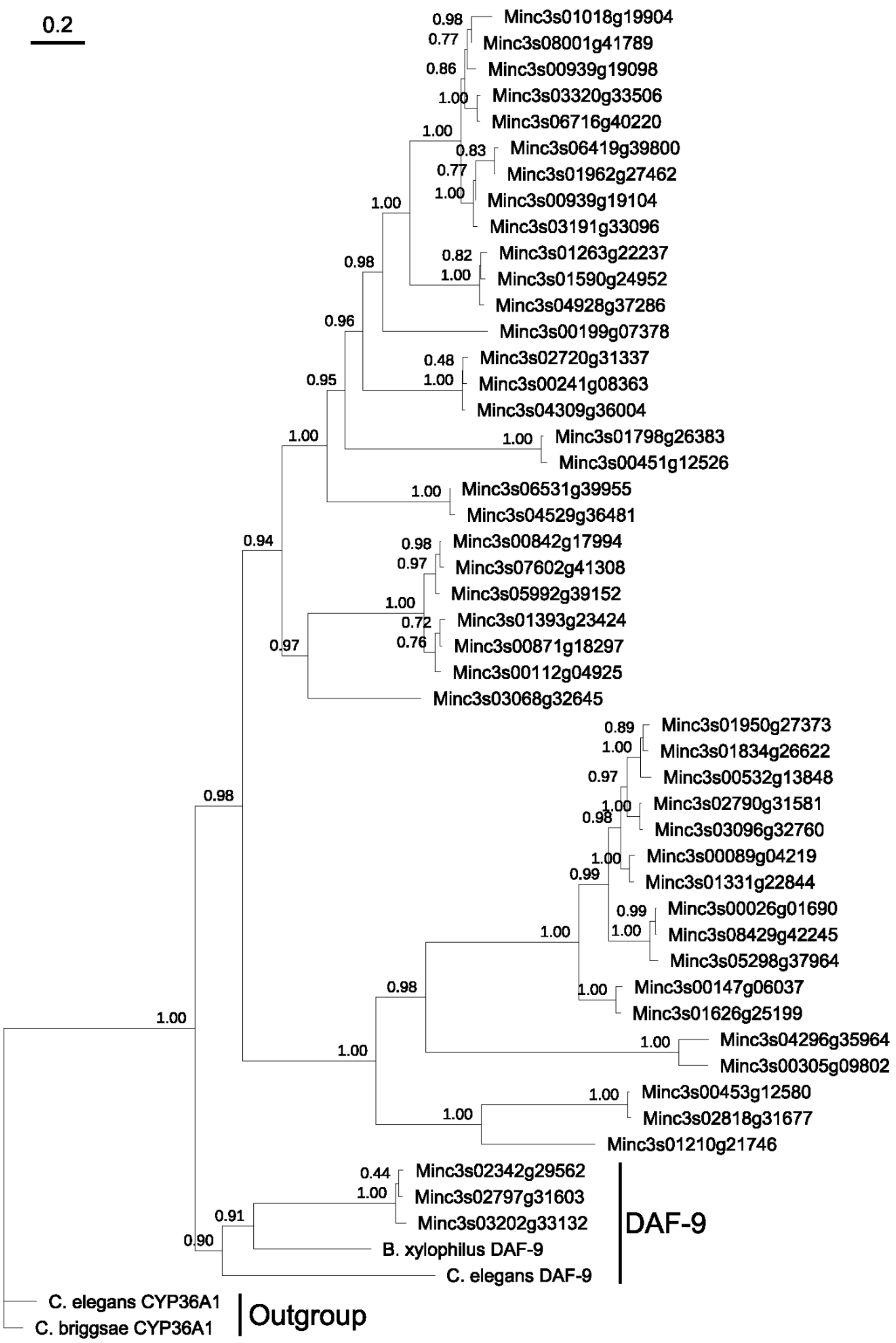


**S7 Fig. DAF-9-*like* nucleotide phylogenetic tree.** Branch length is proportional to genetic change, tree scale shows genetic change per length unit, numbers on the branches indicate bootstrap values, clade label "DAF-9" comprises DAF-9 reference sequences and the 3 final candidates to *M. incognita* DAF-9.

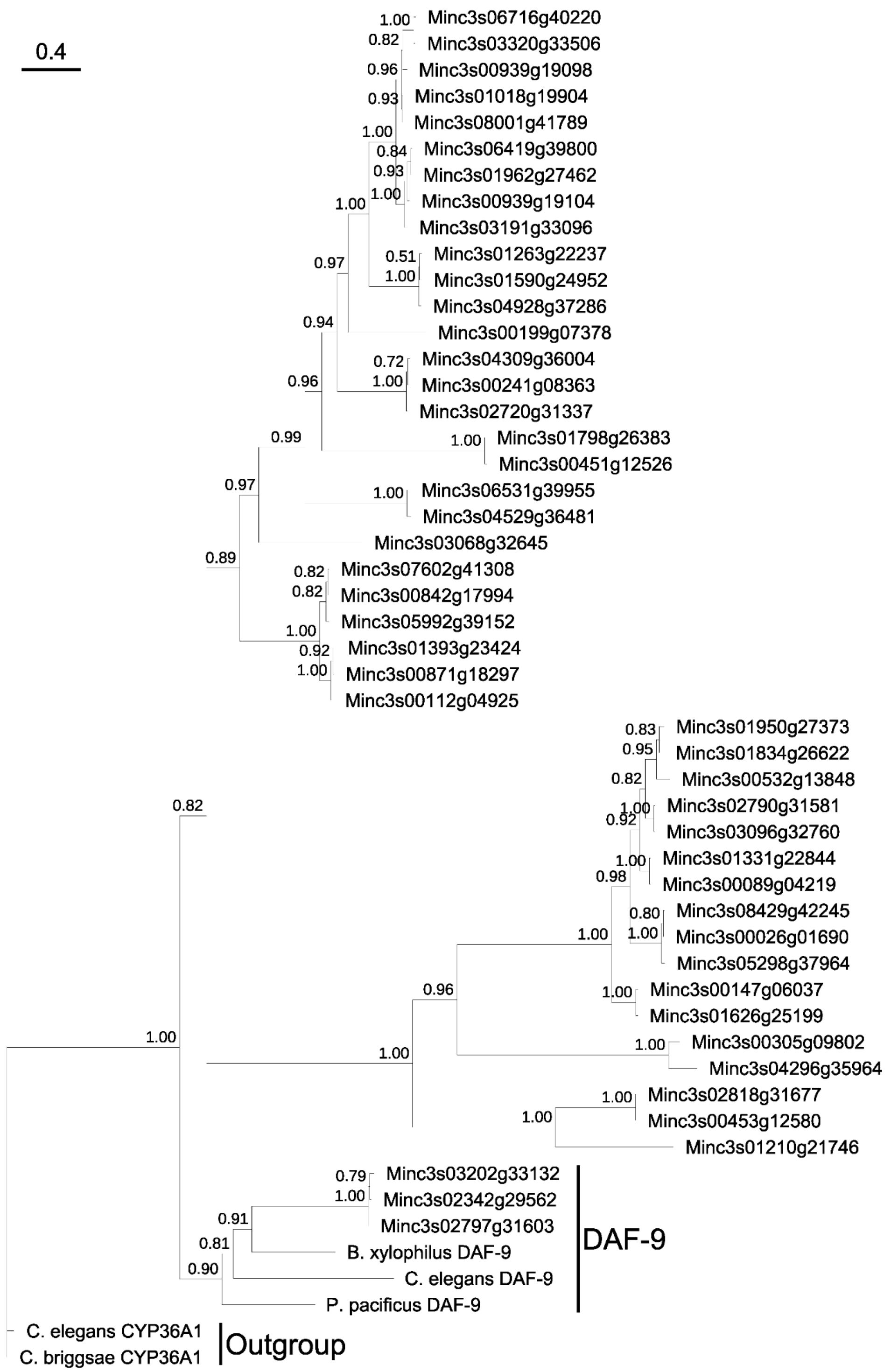


**S8 Fig. DAF-9-*like* protein phylogenetic tree.** Branch length is proportional to genetic change, tree scale shows genetic change per length unit, numbers on the branches indicate bootstrap values, clade label "DAF-9" comprises DAF-9 reference sequences and the 3 final candidates to *M. incognita* DAF-9.

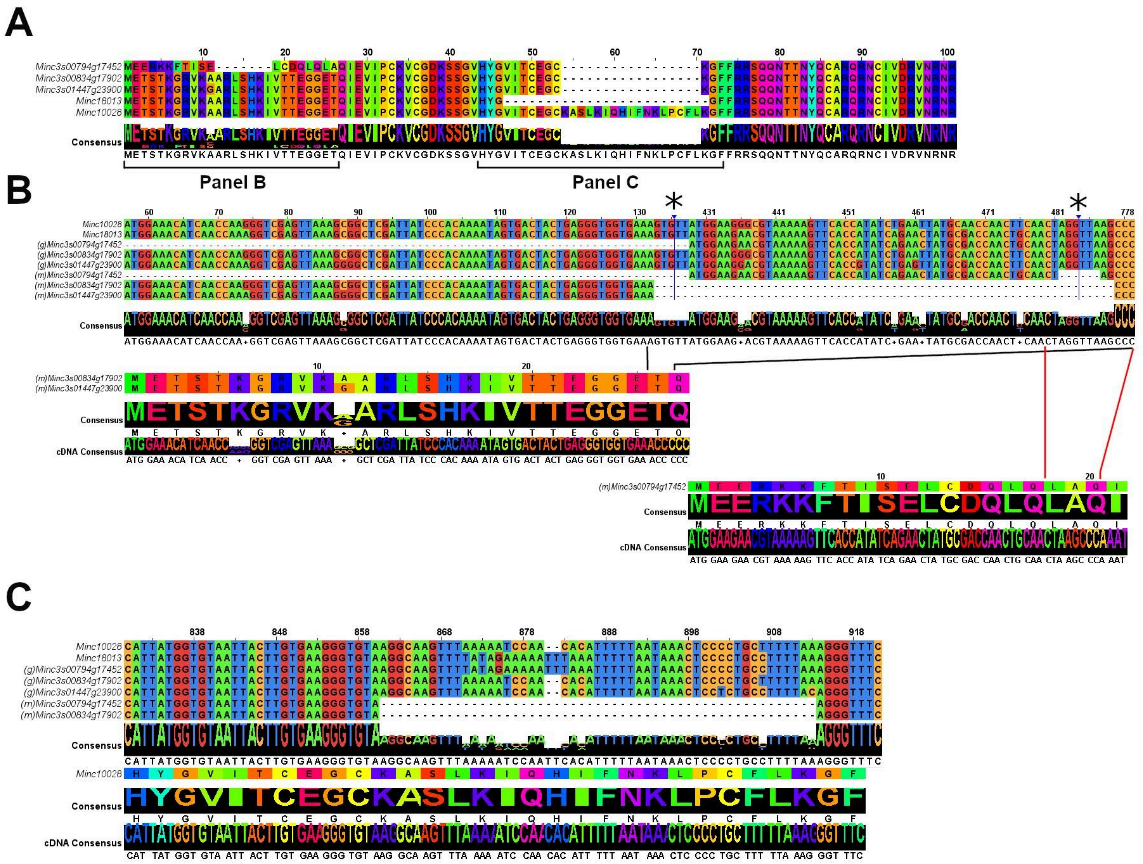


**S9 Fig. Protein, mRNA and genomic M. incognita DAF-12 sequence alignments and logos.** Asterisks above MSAs indicate omitted regions. (A) Minc18013 and Minc10028 are hypothetical DAF-12 protein sequences identified in the *M. incognita* old genome version. Protein sequences beginning with "Minc3s" are the 3 DAF-12 candidates identified in this study. Brackets point out the translation of the genomic sequences in panels B and C. (B) Minc10028 and Minc18013 are the genomic sequences of the hypothetical DAF-12 sequences in the *M. incognita* old genome version. Sequences beginning with "(g)" and "(m)" are the genomic and mRNA sequences of the 3 DAF-12 candidates respectively. Aligned below the nucleotide MSA are the translations of the 3 mRNA sequences. Black and red lines indicate genomic regions skipped in the translation. (C) Sequences are the same as in (B). Below the alignment is the respective protein sequence of Minc10028 that includes the translation of an intronic segment.

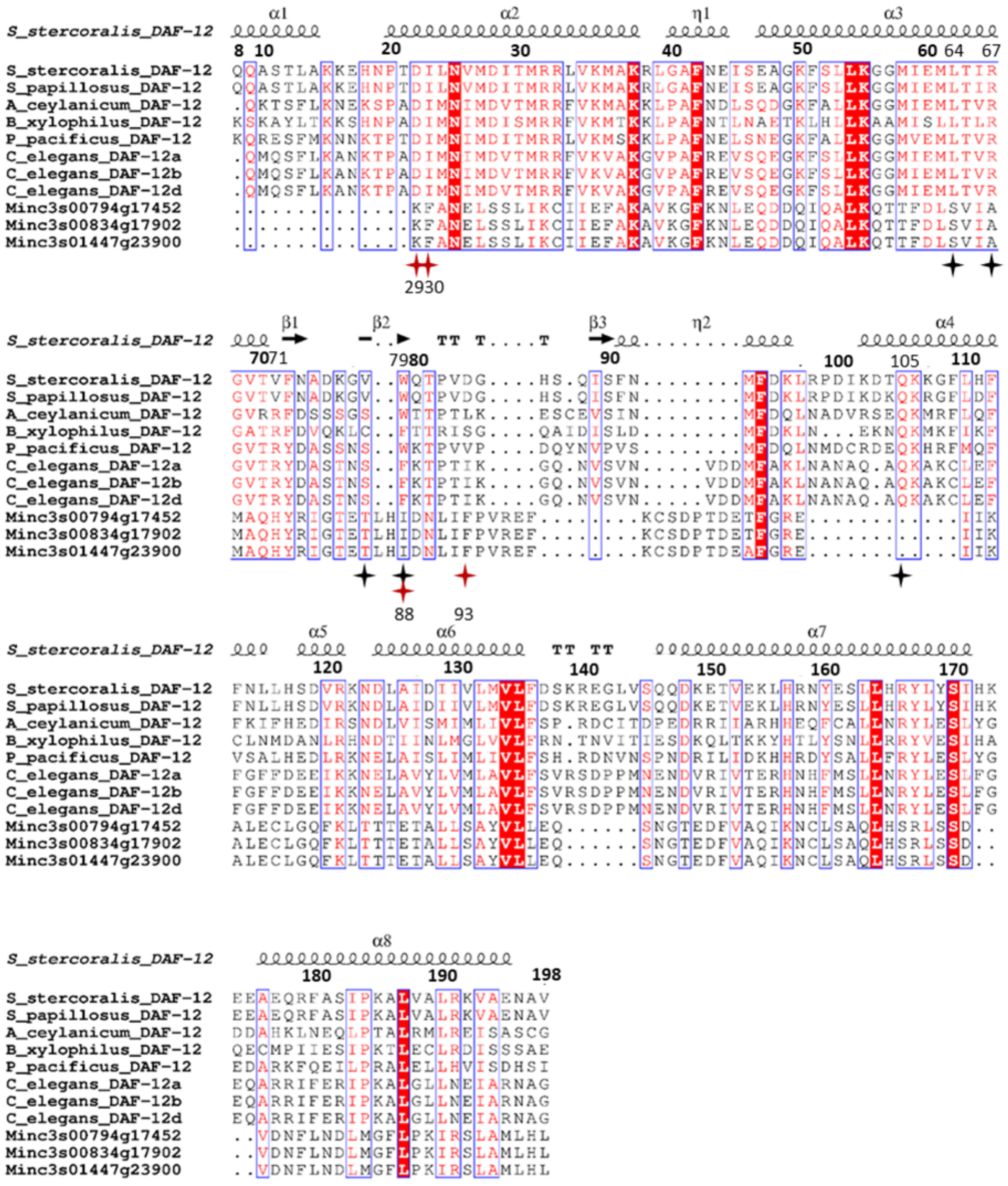


**S10 Fig. DAF-12 protein sequences MSA. Protein structure mapped to MSA is DAF-12$_{Sster}$.** Dots represent gaps, perfectly conserved columns are highlighted in red, highly conserved columns have letters in red, highly conserved regions are marked in blue frames, black stars highlight key amino acids in the *S. stercoralis* hormone receptor binding site that are important to form bonds with ligands, red stars and numbers beneath them highlight key amino acids in the *M. incognita* model binding site (and their positions in the unaligned sequence) that are important to form bonds with ligands. For secondary structure squiggles represent helices (α and η are α-helices and 310-helices respectively), arrows are β-sheets and TT stands for β-turn. The plot was generated with ESPript[44].

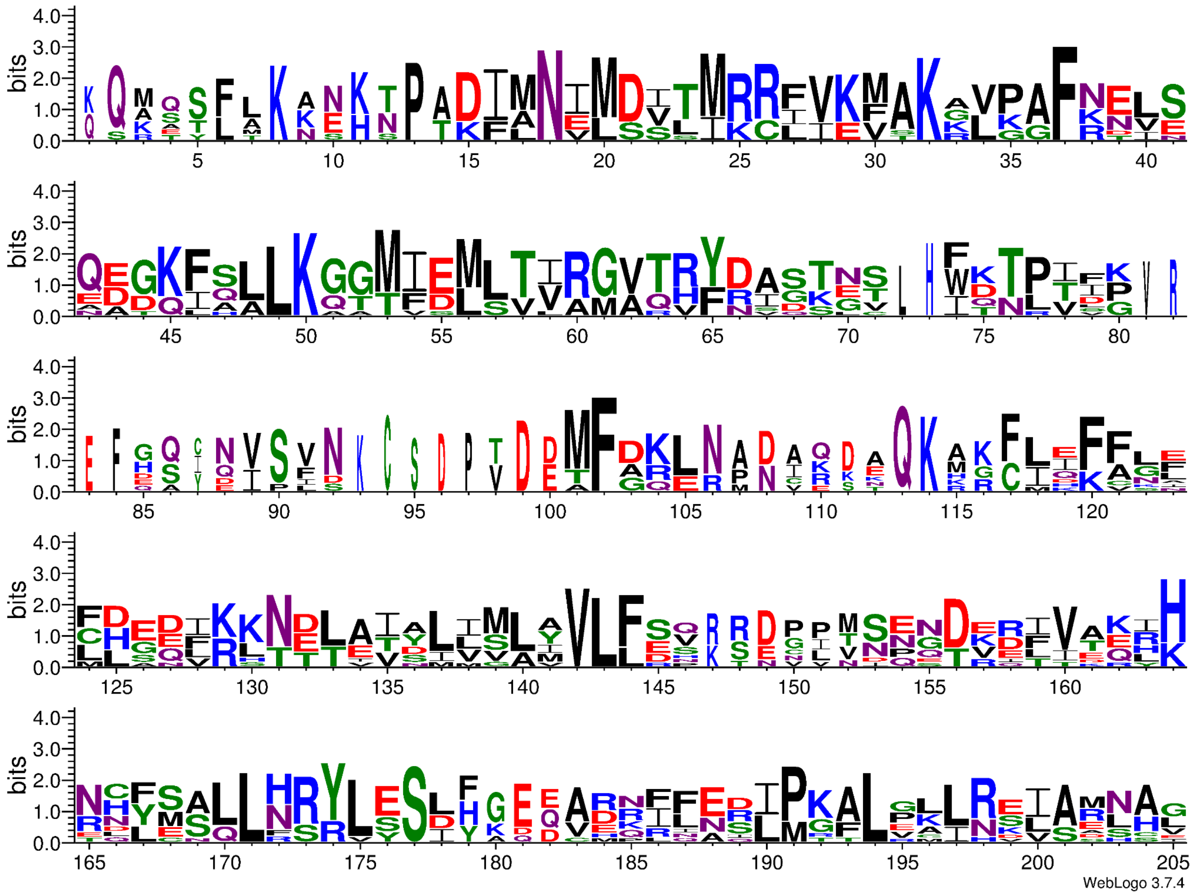

**S11 Fig. DAF-12 protein sequences logo.** Letter height indicates residue conservation in the correspondent column in the MSA, while width indicates gap presence.

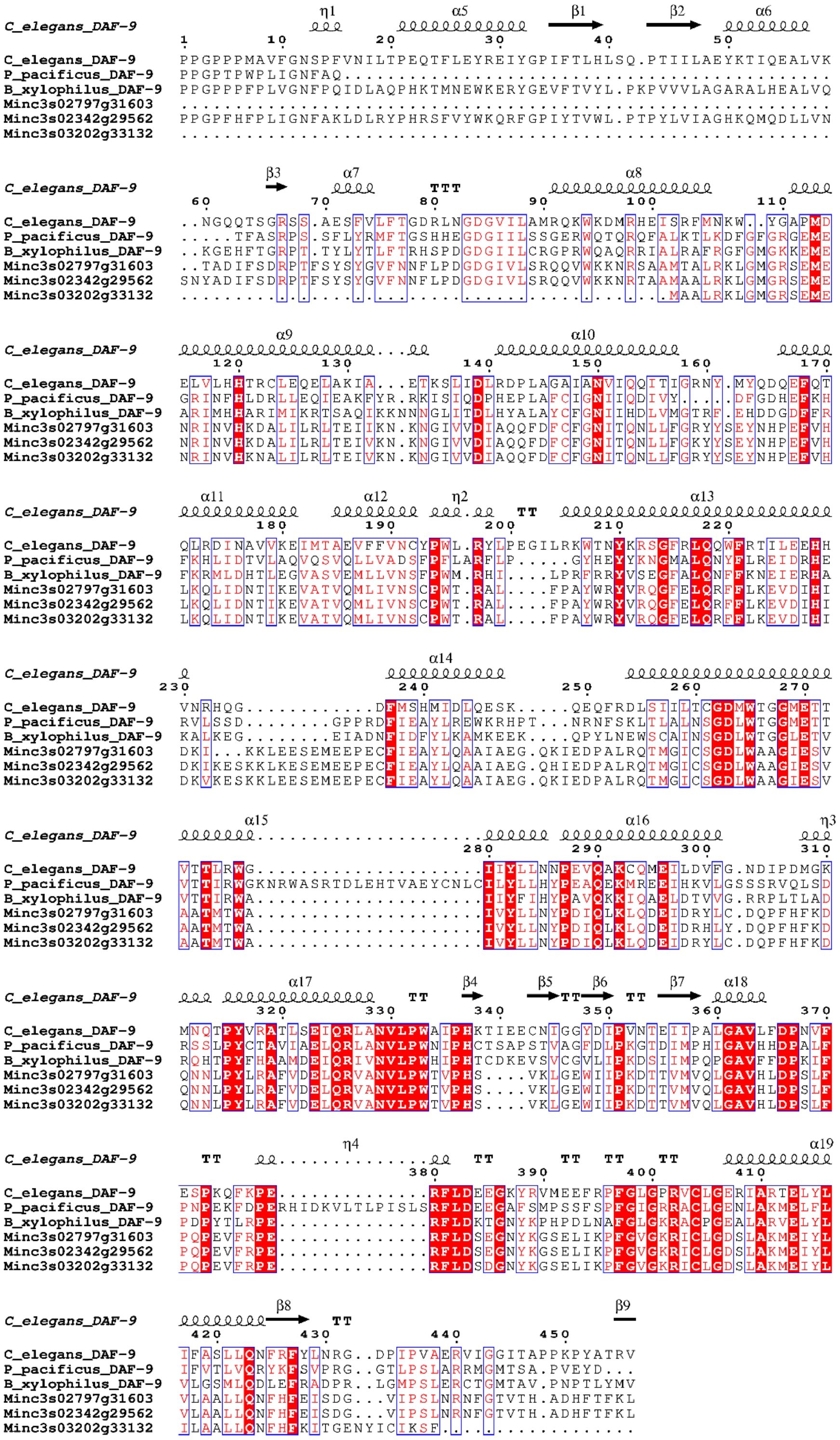

**S12 Fig. DAF-9 protein sequences MSA. Protein structure mapped to MSA is *C. elegans* DAF-9 AlphaFold prediction for Uniprot sequence H2KYS3.** Dots represent gaps, perfectly conserved columns are highlighted in red, highly conserved columns have letters in red, highly conserved regions are marked in blue frames. For secondary structure squiggles represent helices (α and η are α-helices and 310-helices respectively), arrows are β-sheets and TT stands for β-turn. The plot was generated with ESPript[44].

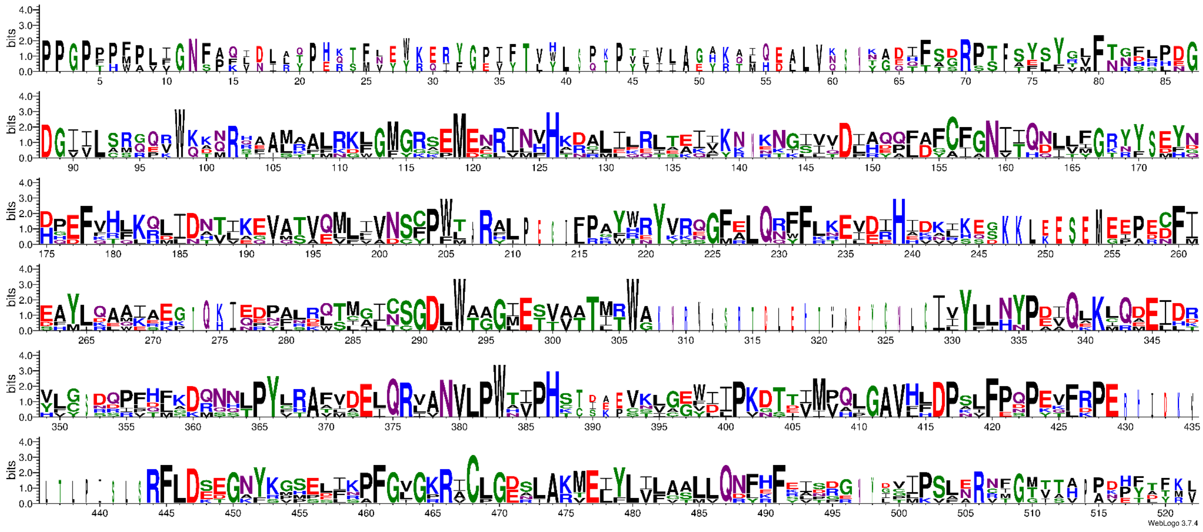

**S13 Fig. DAF-9 protein sequences logo.** Letter height indicates residue conservation in the correspondent column in the MSA, while width indicates gap presence.

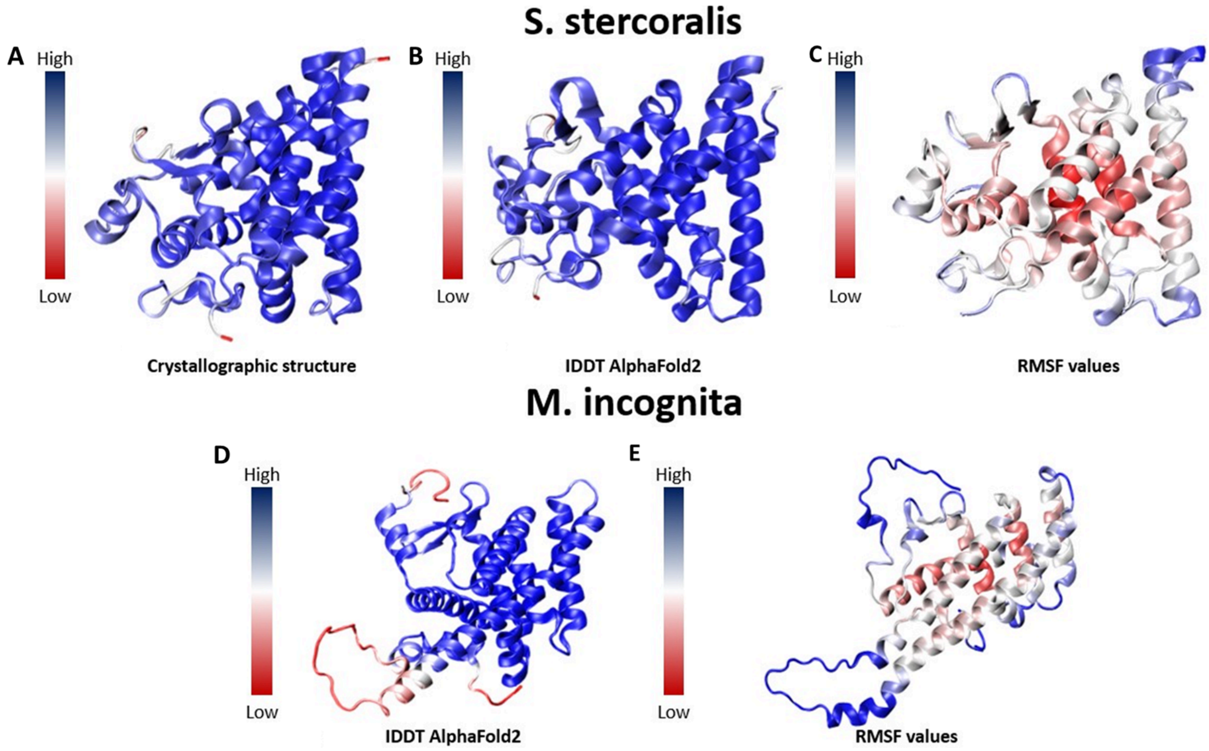


**S14 Fig. Quality of models.** (A) Crystallographic structure of DAF-$12_{Sster}$ colored by BFactor where red parts highlight unstable regions while blue ones represent highly stable regions. (B) AlphaFold2 3D model of DAF-$12_{Sster}$. In Blue high local model quality IDDT regions and in red regions with low IDDT values. (C) AlphaFold2 3D MD-refined model of DAF-$12_{Sster}$ colored by RMSF values show in red those regions that had the most mobility during the simulations while blue ones show those regions that remain more stable. (D) AlphaFold2 3D model of DAF-$12_{Minc}$. In Blue high local model quality IDDT regions and in red regions with low IDDT values. (E) AlphaFold2 3D MD-refined model of DAF-$12_{Minc}$ colored by RMSF values show in red those regions that had the most mobility during the simulations while blue ones show those regions that remain more stable.

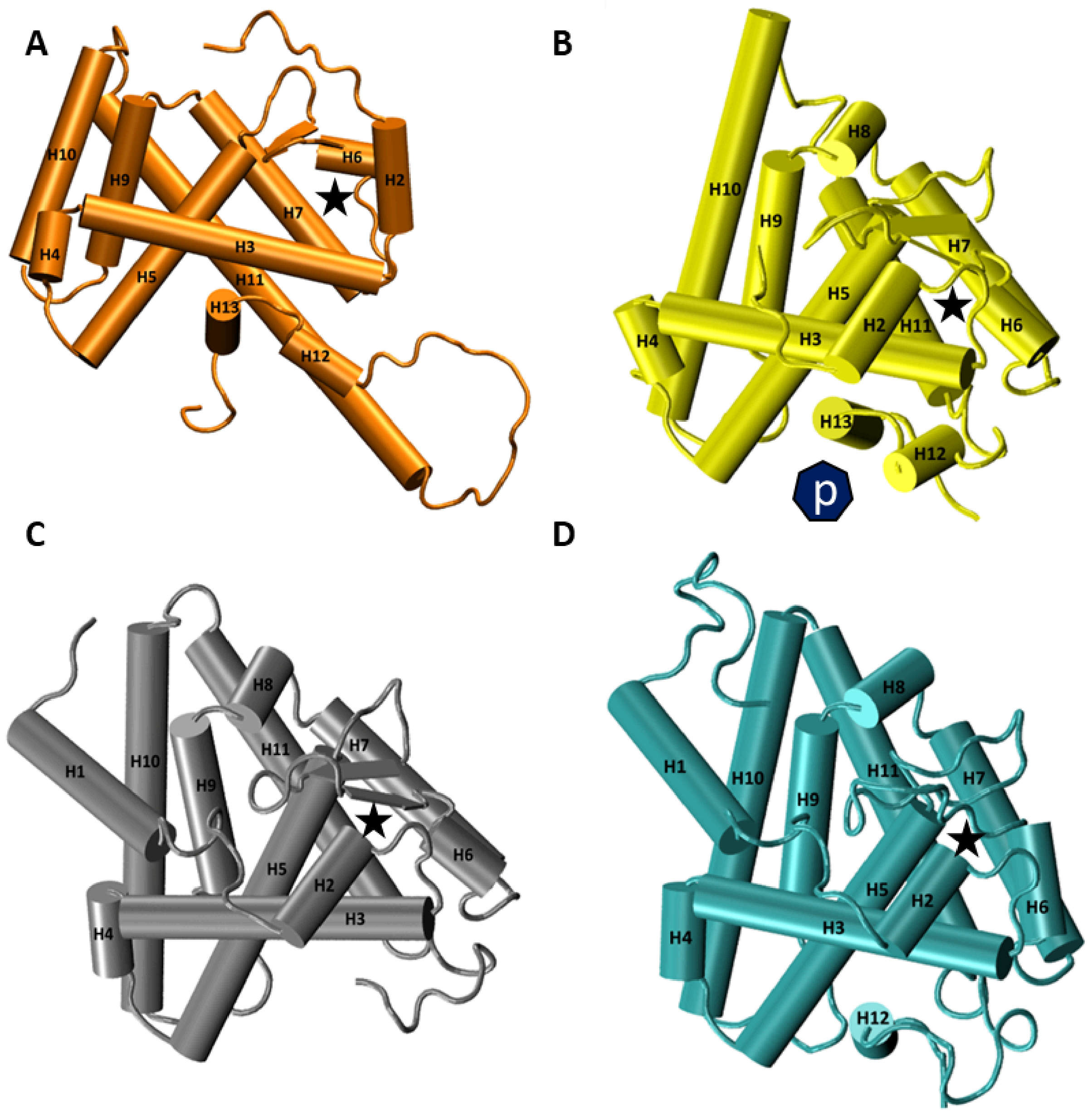


**S15 Fig. Secondary structure comparison.** DAF-12 receptor showed in cartoon style of DAF-12$_{Minc}$ (A), DAF-12$_{Sster}$ with signal peptide as a P in black heptagon, DAF-12$_{Ace}$ (C) and DAF-12$_{Cele}$ (D). Labels in alpha helices indicate its number. In all models the star highlights the binding site of the protein.

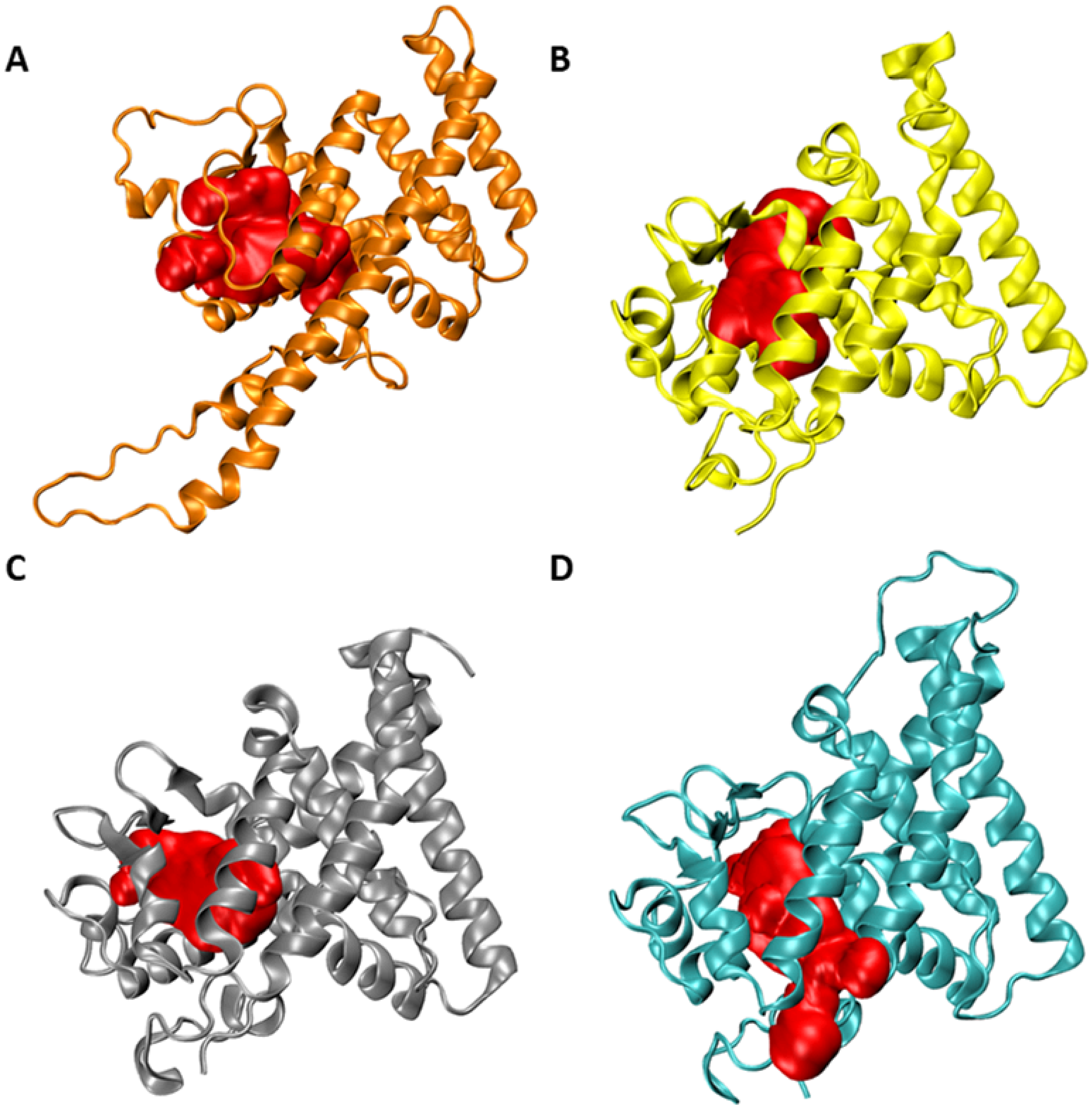


**S16 Fig. Pocket search.** Protein structures are shown in cartoon style while red bodies represent the Pocket found in DAF-12$_{Cele}$ *(A),* DAF-12$_{Minc}$ (B), DAF-12$_{Sster}$ (C) and DAF-12$_{Ace}$ *(D)*.

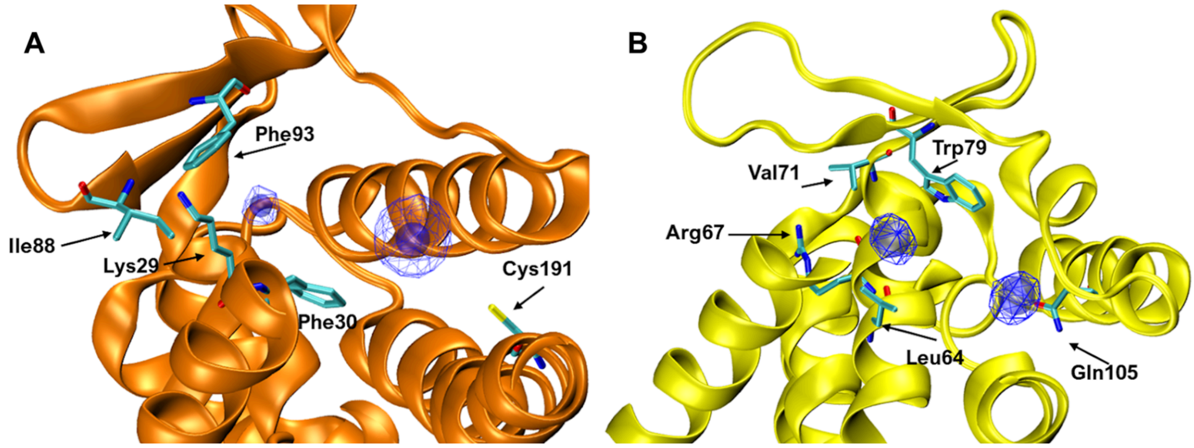


**S17 Fig. Co-solvent molecular dynamics reveals important amino acid interactions in DAF-12.** The structures for the DAF-12 hormone receptor for *M. incognita* and *S. stercoralis* are presented in panel A and B as cartoon representations colored as orange and yellow respectively. Key amino acids are represented as sticks, colored by atoms (red, blue, yellow and cyan for oxygen, nitrogen, sulfur and carbon respectively). The high occupancy regions are shown as blue lines and balls.

**S1 Table. Secondary structure proportions.** Secondary structure proportions among the proteins.

| **Structure** | $DAF\text{-}12_{Minc}$ | | $DAF\text{-}12_{Sster}$ | | $DAF\text{-}12_{Acey}$ | | $DAF\text{-}12_{Cele}$ | |
|---|---|---|---|---|---|---|---|---|
| **Alpha helix** | **11** | **39.3%** | **12** | **38.7%** | **13** | **41.9%** | **12** | **41.4%** |
| **Loop** | **14** | **50.0%** | **16** | **51.6%** | **15** | **48.4%** | **14** | **48.3%** |
| **Beta sheet** | **3** | **10.7%** | **3** | **9.7%** | **3** | **9.7%** | **3** | **10.3%** |

**S2 Table. Binding site characterization.** Most relevant parameters estimated with Fpocket software for each refined structure.

| Protein | Model | Pocket Score | Drug score | Hydrophobicity score | Polarity score | Volume A^3 |
|---|---|---|---|---|---|---|
| **DAF-12$_{Sster}$** | **Unrefined model** | 30.859 | 0.759 | 39.133 | 7 | 457.933 |
| | **Refined model** | 34.621 | 0.903 | 35.824 | 13 | 1655.032 |
| **DAF-12$_{Ace}$** | **Unrefined models** | 38.500 | 0.776 | 32.250 | 11 | 831.504 |
| | **Refined model** | 37.356 | 0.762 | 27.345 | 11 | 1076.730 |
| **DAF-12$_{Cele}$** | **Unrefined models** | 37.368 | 0.774 | 28.524 | 9 | 740.727 |
| | **Refined model** | 27.400 | 0.511 | 27.700 | 26 | 2158.185 |
| **DAF-12$_{Minc}$** | **Unrefined model** | 40.582 | 0.730 | 24.364 | 17 | 1206.552 |
| | **Refined model** | 43.495 | 0.788 | 29.816 | 23 | 2236.824 |

**S1 File. Fasta file containing the 3 *M. incognita* DAF-12 protein candidates.**

**S2 File. MSA used for DAF-12 nucleotide identity matrix.**

**S3 File. MSA used for DAF-12 protein identity matrix.**

**S4 File. MSA used for DAF-9 nucleotide identity matrix.**

**S5 File. MSA used for DAF-9 protein identity matrix.**

**S6 File. DAF-12-*like* proteins filtered from the proteome through data mining.**

**S7 File. DAF-9-*like* proteins filtered from the proteome through data mining.**
